# Exploring the Plant Aquaporin Solute Transport Network: Functional characterisation of *Nicotiana tabacum* PIP, TIP and NIP isoforms

**DOI:** 10.1101/2021.03.26.437249

**Authors:** Annamaria De Rosa, Rose Zhang, Caitlin Byrt, John R Evans, Michael Groszmann

**Affiliations:** ARC Centre of Excellence for Translational Photosynthesis, Research School of Biology, Australian National University, 134 Linnaeus Way, Canberra, ACT 2601, Australia; Plant Science Division, Research School of Biology, Australian National University, 134 Linnaeus Way, Canberra, ACT 2601, Australia

## Abstract

Aquaporins (AQPs) are multifunctional membrane proteins which have greatly diversified in number and function in the plant Kingdom. In plants, AQPs have evolved to comprise a dynamic solute transport network occurring in all tissues and facilitating transport of water and vital solutes across various cellular membranes. Plant AQPs are involved in a multitude of plant physiological processes, however a better understanding is required of AQP structure-function relationships, multifunctionality and cell membrane localisation in order to begin to describe putative functional roles for the numerous plant AQP gene isoforms. Using an integrated approach, we characterised nine diverse *Nicotiana tabacum* (tobacco) aquaporins, spanning the 3 largest AQP subfamilies (PIP, TIP, and NIP) and with varied gene expression profiles. High-throughput yeast-based functional screens identified novel candidates for water, hydrogen peroxide (H_2_O_2_), boric acid (BA) and urea transport across the 3 AQP subfamilies. Using GFP translational fusions, AQPs observed *in planta* were localised to the plasma membrane, tonoplast and endoplasmic reticulum. AlphaFold protein models illustrated differences in pore shape and size across subfamilies. Our analysis supports the importance of functional data for deciphering unknown AQP structure-function relationships and uncovering novel candidates for *in planta* solute transport.

## Introduction

Improving our understanding of solute transport in plants could enhance our potential to harness plant biology to deliver novel plant-based biotechnological innovations. *Nicotiana tabacum* (tobacco) is a commonly used model system, and more broadly has renewed commercial applications including its uses as a bio-factory for production of biofuels and plant-made pharmaceuticals (Vanhercke et al. 2014; Tusé et al. 2014; Ma et al. 2015). Furthermore, its relatedness to horticulturally important crops such as tomato, potato, eggplant and capsicum, makes tobacco a favourable study species for translating key findings into crops (De Rosa et al. 2020).

Aquaporins (AQPs) constitute a major family of integral membrane channel proteins found across all kingdoms of life (Abascal et al. 2014), becoming most diversified in number and subfamilies in higher plants (Groszmann et al. 2017; Abascal et al. 2014). Plant AQPs play a vital role in diverse physiological processes, including water relations, growth and development, stress responses, and photosynthesis (Hachez et al. 2006; Groszmann et al. 2017; Chaumont and Tyerman 2017; Zhang et al. 2023). This range in cellular functions reflects a capability for transporting a wide variety of substrates including water, nitrogen compounds (e.g. ammonia, urea and nitrate), gases (e.g. carbon dioxide, oxygen), hydrogen peroxide, metalloids (e.g. boron, silicon), and ions (Gomes et al. 2009; Pommerrenig et al. 2015; Hove and Bhave 2011; Choi and Roberts 2007; Zwiazek et al. 2017; Byrt et al. 2017; Bienert et al. 2013; Liu et al. 2020). AQPs are part of a dynamic network for transport of water and vital solutes in all plant tissue types and across several cellular membranes (plasma membrane, tonoplast, chloroplast envelope, mitochondria, endoplasmic reticulum). However, functional characterisation of AQPs for multiple substrates is quite limited and elucidating substrate transport profiles of various isoforms is a key step towards understanding the diverse potential AQP functional roles in plants and further investigating the many regulatory factors which precisely control AQP cell-to-cell and cellular solute transport e.g. AQP-AQP interactions, AQP-protein interactions and cellular signals (Roche and Törnroth-Horsefield 2017; Bellati et al. 2016; Fox et al. 2017).

Aquaporins assemble as tetrameric complexes, with each monomer forming a functional pore created by six membrane spanning helices, five connecting loops and two shorter helices. Formation of tetrameric AQP complexes creates a central fifth and functioning pore (Pommerrenig et al. 2015; Kirscht et al. 2016; Törnroth-Horsefield et al. 2006). Although the gross tertiary structure of AQP is highly conserved across organisms, slight deviations in structural and functional characteristics between isoforms contribute to differences in their transport selectivity. Such characteristics include pore dimensions (pore diameter and overall morphology), chemical properties and flexibilities of pore-lining residues, and specific configurations of residues at key constriction points (Luang and Hrmova 2017). Higher plant AQPs divide into five phylogenetically distinct sub-families, namely the Plasma membrane Intrinsic Proteins (PIPs), Tonoplast Intrinsic Proteins (TIPs), Small basic Intrinsic Proteins (SIPs), Nodulin 26-like Intrinsic Proteins (NIPs), and X Intrinsic Proteins (XIPs) (Danielson and Johanson 2008; Johanson and Gustavsson 2002; Kaldenhoff and Fischer 2006). Within each of these sub-families, there can be diversity in permeating substrate selectivity and specific organelle membrane integration (Maurel et al. 2008). Subfamily-specific substrate specificities in plant AQPs have been attributed to diversity in the aromatic arginine (ar/R) selectivity filter (SF) which forms the first constriction site towards the extracellular side of the pore (Hove and Bhave 2011). Variation in this site largely determines the substrates able to permeate across the membrane through the AQP (Hove and Bhave 2011; Sui et al. 2001). Dual Asn-Pro-Ala (NPA) motifs located at the centre of the pore act as a second constriction, with variation in residue composition contributing to selectivity for substrates such as ammonia, boric acid (BA) and urea (Wu and Beitz 2007; Hove and Bhave 2011).

Teasing apart the complexities of aquaporin biology *in planta* is difficult. Plants have many AQP isoforms, with tobacco in particular arising from a recent hybridization and its allotetraploid genome containing 84 AQP genes (De Rosa et al. 2020; Ahmed et al. 2020; Groszmann et al. 2021). Such large AQP families have the potential for redundancy of function under certain environmental conditions (Abascal et al. 2014; Fox et al. 2017). As such, modification of aquaporin expression within the plant, via over expression or down regulation of a specific AQP, may also affect the expression of closely related isoforms (Bi et al. 2015; Kaldenhoff et al. 2007).

Functional characterisation of plant AQPs is often accomplished through heterologous expression systems, such as yeast (Kaldenhoff et al. 2007), allowing for identification of putative solute transport for an AQP homo-tetramer in isolation of other isoforms and confounding external regulatory mechanisms (Groszmann et al. 2023). We used high-throughput micro cultivation-based yeast assays (described in Groszmann et al. 2023) to screen tobacco AQPs spanning the 3 largest AQP subfamilies, namely the PIPs, TIPs and NIPs. These AQPs isoforms were chosen based on homology to already characterised AQPs in other species or on gene expression characteristics within the plant which implied diverse functional roles. Three PIP1 genes were chosen: NtPIP1;5s (NtAQP1), NtPIP1;1t and NtPIP1;3t, each sharing more than 90% homology in gene sequence. NtPIP1;5s (NtAQP1) is an established CO_2_-permeable AQP isoform, that enhanced CO_2_ diffusion and photosynthetic efficiency in planta (Uehlein et al. 2003; Flexas et al. 2006). Two PIP2 genes, NtPIP2;4s and NtPIP2;5t, were chosen as representative isoforms from distinct phylogenetic sub-clades within the PIP2 phylogeny. From the TIPs, NtTIP1;1s was chosen as a gene highly expressed throughout the plant (De Rosa et al. 2020) and potentially a candidate permeable to a range of solutes, while NtTIP2;5t is predominantly expressed in roots (De Rosa et al. 2020) with high homology to AtTIP2;1, an established transporter of nitrogen compounds (urea and ammonium) (Loqué et al. 2005; Liu et al. 2003; Kirscht et al. 2016). Representative isoforms from two distinct subclasses within the NIP subfamily were selected. NtNIP5;1s in the NIP II sub-class and homologous to the boron-permeable AtNIP5;1 (Takano et al. 2006; Pommerrenig et al. 2015). Whereas, NtNIP2;1s belongs to the NIP III subclass, hypothesised to transport a diverse range of metalloid compounds including the important micronutrients boron and silicon.

We functionally screened these 9 NtAQPs for transport of water and three neutral solutes essential for plant growth: hydrogen peroxide (H_2_O_2_), BA and urea. Sub-cellular localisation of each AQP was visualised *in planta* with an Arabidopsis expression system. Putative substrate transport profiles and sub-cellular localisation data were integrated with gene expression data and AlphaFold models to characterise these tobacco AQPs. Functional screening of these diverse tobacco AQPs identified novel and multifunctional water, H_2_O_2_, urea and BA-permeable candidates, which form part of complex and dynamic cellular solute transport networks.

## Results

### Diverse PIP, TIP and NIP isoforms localised to the yeast plasma membrane

We examined the localisation of 9 diverse NtAQP isoforms. These isoforms span the 3 largest AQP subfamilies, PIP, NIP and TIP, and fall in distinct clades in the NtAQP phylogeny, with the exception of the NtPIP1s (*NtPIP1;1t, NtPIP1;3t* and *NtPIP1;5s*) which are highly homologous gene isoforms (Figure 1) with >90% sequence identity. N-terminal GFP-AQP fusions were used to establish whether these NtAQP proteins localised to the plasma membrane (PM) of yeast (Figure 2A-I), an important feature when using heterologous expression systems for functional screening experiments. Optically thin focal plane images of each yeast construct show the localisation of the GFP signal. These images were further processed using surface profiling and line scans of signal intensity to better assess the distribution of GFP-AQP within the yeast cells. Yeast expressing GFP alone (Figure 2J), showed a uniform signal throughout the cell with the exception of the vacuole. The surface and line signal scans show a relatively equal distribution of intensity, consistent with cytosolic localisation (Figure 2J). The fusion of NtAQPs to GFP resulted in the redistribution of GFP fluorescence to different yeast sub-cellular compartments including; PM, endoplasmic reticulum (ER), and/or the tonoplast (vacuolar membrane). The GFP-NtPIP1 fusions localised to the periphery of the cell and the ER. Although signal intensity in the periphery varied, likely due to co-localisation in peripheral ER, the signal was continuous around the cell, consistent with PM integration (Figure 2 A-C). In addition to localisation in the yeast PM, images for NtPIP1;5s frequently contained bright spots in the periphery of the cell which are characteristic of ER localisation (Figure 2C). The NtPIP2 proteins integrated into the PM, with clearly defined peaks present in the line scans, but GFP signal also localised to the ER and faintly inside the vacuole (Figure 2E-D). NtNIP2;1s localised to the PM and ER, similar to that observed for the NtPIP1s (Figure 2F). NtNIP5;1 weakly localised to the PM and ER, with signal predominantly associated with integration into the tonoplast (Figure 2G). The NtTIPs had strong signals distributed between the ER, tonoplast and notably the PM (Figure 2H-I). Overall, although our yeast localisation images showed variation in brightness and allocation of AQPs to various membranes (PM, tonoplast and ER) or to the vacuole, we could confirm PM integration for all NtAQP constructs tested, enabling further testing of solute transport across the PM in our yeast-based functional screens.

**Figure 1.**
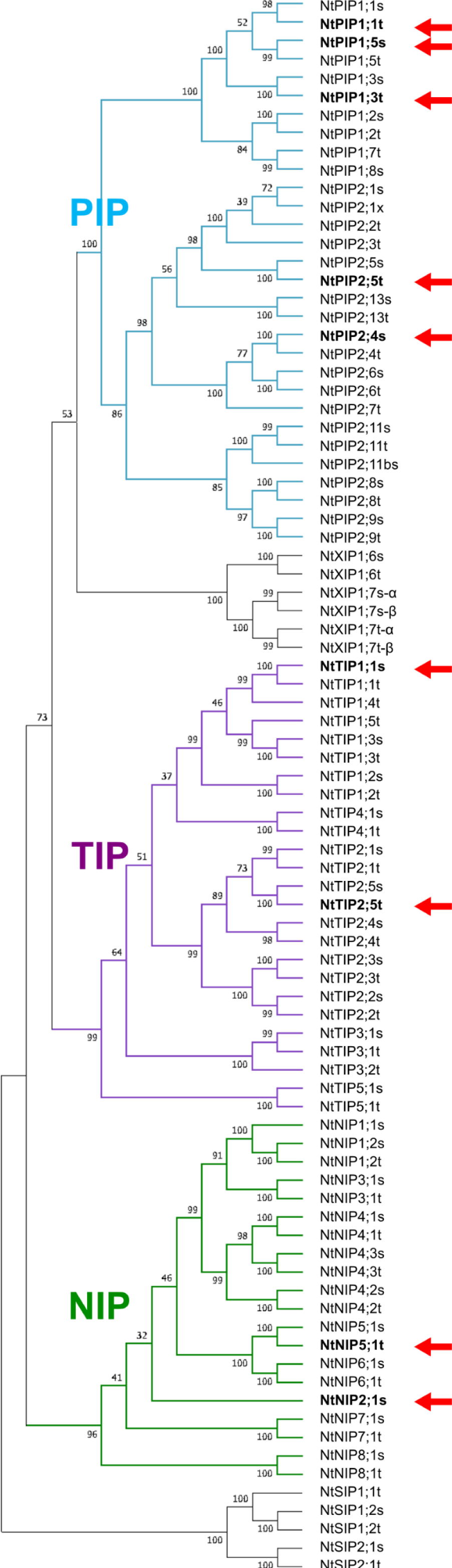
Phylogeny of the NtAQP family, highlighting genes selected for functional characterisation in this study. The phylogenetic tree was generated using the neighbour-joining method from MUSCLE aligned protein sequences. Confidence levels (%) of branch point generated through bootstrapping analysis (n=1000). Red arrows point to NtPIP (NtPIP1;1t, NtPIP1;3t, NtPIP1;5s, NtPIP2;4s, NtPIP2;5t), NtTIP (NtTIP1;1s, NtTIP2;5t) and NtNIP (NtNIP2;1s and NtNIP5;1t) isoforms functionally characterised in this study.

**Figure 2.**
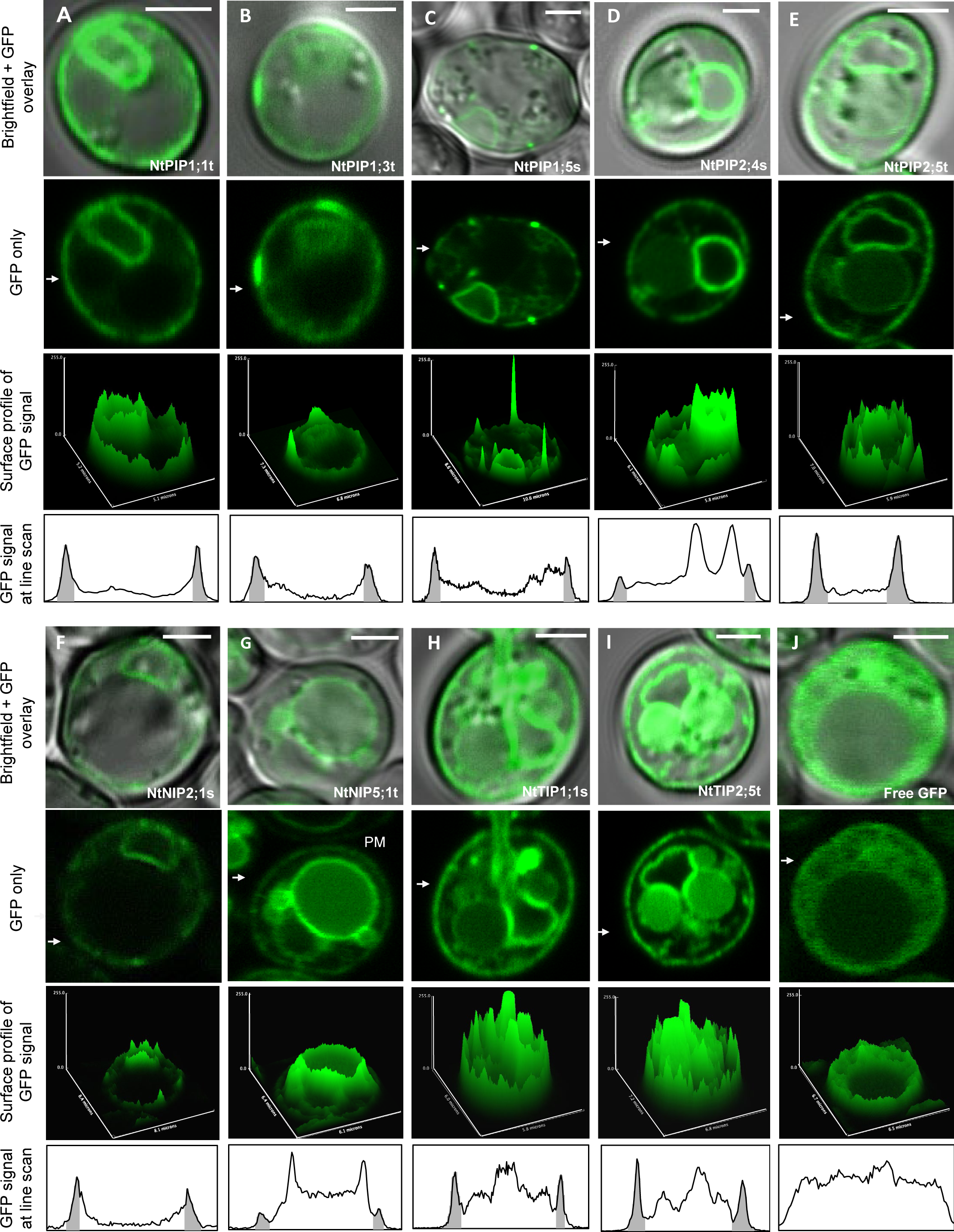
Subcellular localisation of GFP tagged aquaporins expressed in yeast. Confocal microscopy images of yeast expressing GFP::NtAQP fusions of *A*. NtPIP1;1t, *B*. NtPIP1;3t, *C*. NtPIP1;5s, *D*. NtPIP2;4s, *E*. NtPIP2;5t, *F*. NtNIP2;1s, *G*. NtNIP5;1t, *H*. NtTIP1;1s, *I*. NtTIP2;5t and *J*. Free GFP localisation. For each construct we report a Brightfield + GFP overlay image of a yeast cell; a GFP only image; a surface plot profile of GFP signal intensity at the imaged focal plane; and a line scan of signal intensity traversing the cell (indicated by white arrow in GFP only image). Grey shading in GFP signal line scan corresponds to regions which align with the plasma membrane (PM). PM, endoplasmic reticulum (ER) and vacuole (V) are labelled. Scale bar 2μm.

### Water permeability “Freeze-thaw” assay

High-throughput yeast assays were used to assess growth/survivorship phenotypes of *aqy1 aqy2* yeast exposed to freezing-thawing treatments. The *aqy1 aqy2* yeast strain lacks its native water transporting AQPs (Tanghe et al. 2002). Ectopic expression of water-permeable NtAQP candidates reduced cell rupture upon freezing, conferring a growth advantage compared to yeast expressing Empty vector control (Groszmann et al. 2023). *aqy1 aqy2* yeast expressing the Empty vector did not survive exposure to two freeze-thaw cycles and failed to grow after treatments (Figure 3A). By contrast, the growth of *aqy1 aqy2* yeast expressing a Freeze-thaw tolerant AQP, *NtPIP2;4s*, subjected to two freeze-thaw cycles was only slightly delayed compared to untreated yeast (Figure 3B). Substantial differences in growth were observed between treated and untreated yeast expressing either a particular NtAQP or empty vector (Figure 3C). Following exposure to two freeze-thaw cycles, the *aqy1 aqy2* yeast expressing the empty vector had only 2% growth relative to the untreated empty vector culture (Figure 3C). NtPIP2;4s and NtPIP2;5t expression resulted in the greatest yeast survival following the freeze-thaw treatments, achieving 70% of the untreated growth (Figure 3C). NtTIP1;1s and NtTIP2;5t yeast grew to 62% and 30%, respectively, relative to untreated controls. Thus, expression of four of the nine NtAQPs tested enabled yeast to survive two freeze-thaw cycles, indicative of water transport across the PM.

**Figure 3.**
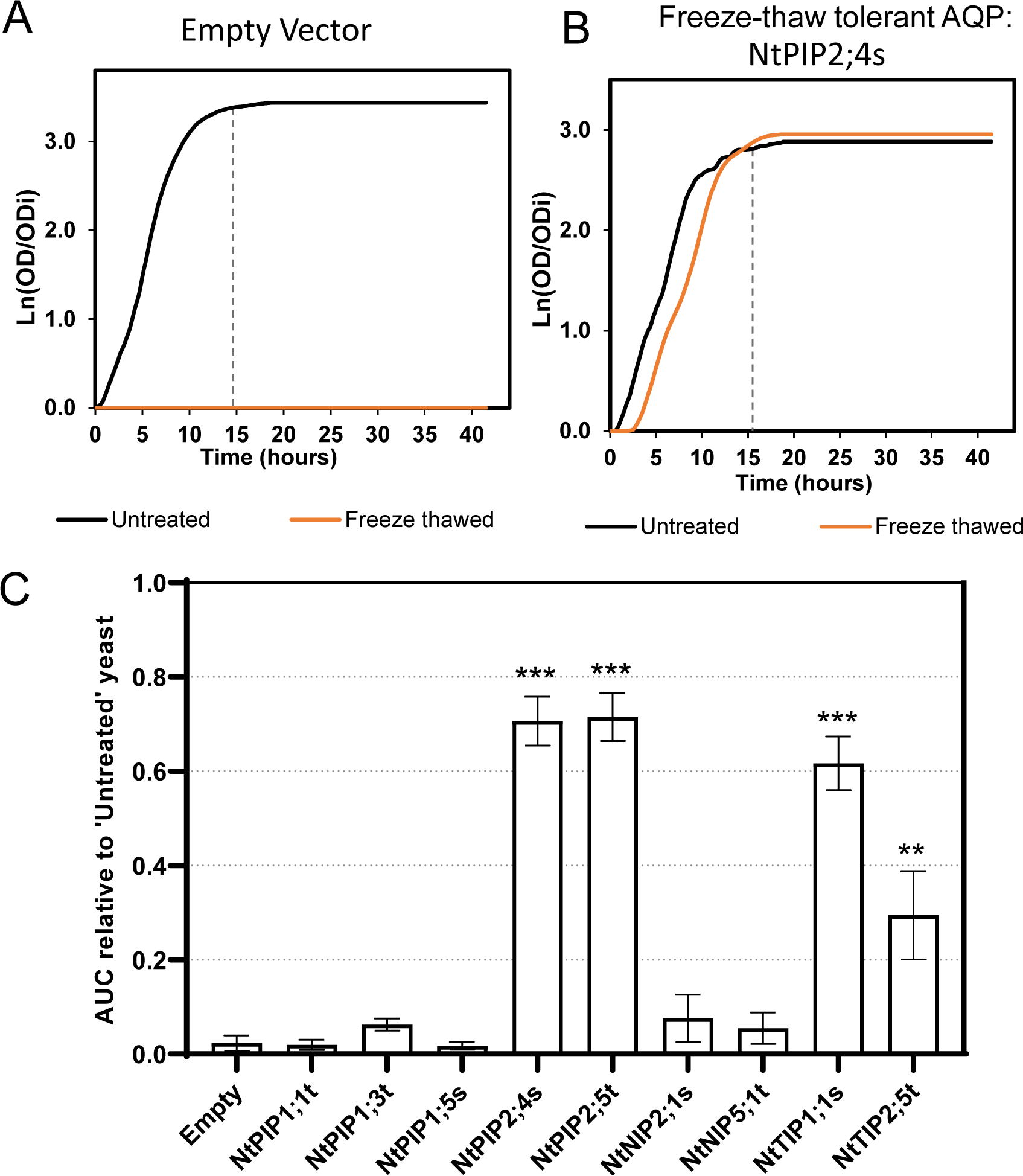
‘Freeze-thaw’ assay testing for NtAQP water transport. Yeast growth curves, Ln(OD/OD_i_) vs. time, of *aqy1 aqy2* yeast expressing **A.** Empty vector control or **B.** a ‘Freeze-thaw tolerant’ AQP (NtPIP2;4s), exposed to freeze-thaw treatments. Growth was assessed from the area under the curves (AUC) until the vertical dashed lines *representing Phi ф*. **C.** Yeast culture growth following the freeze–thaw treatment (AUC relative to untreated yeast control) for *aqy1 aqy2* yeast expressing an Empty vector or one of the 9 NtAQPs. Asterisks denotes significantly greater growth following the freeze – thaw treatment compared against Empty vector from an ANOVA: **p<0.01 and ***p<0.001, N=6, Error bars = SE.

### H_2_O_2_ toxicity assay

Toxicity-based assays were used to compare H_2_O_2_-sensitivity of skn7 (ROS hypersensitive) yeast expressing the NtAQP isoforms and Empty vector control. Dose-dependent differences in growth curves of yeast expressing the empty vector (Figure 4A) or a “H_2_O_2_-sensitive” AQP candidate (Figure 4B) were observed when exposed to increasing H_2_O_2_ concentrations. Yeast expressing the empty vector and exposed to 0.25mM or 0.5mM H_2_O_2_ treatments, had no significant reduction on growth relative to untreated yeast, while 1mM H_2_O_2_ treatment caused a 37% decrease in growth relative to the untreated control (Figure 4C). By contrast, growth was dramatically reduced in the presence of 0.25mM H_2_O_2_ for yeast expressing NtPIP2;4s (growth reductions were 66%, 86% and 80% at 0.25mM, 0.5mM and 1mM H_2_O_2_, respectively, Figure 4C). Five NtAQPs increased yeast sensitivity to H_2_O_2_ exposure compared to empty vector, consistent with these AQPs being classified as H_2_O_2_ permeable candidates: NtPIP2;4s, NtPIP2;5t, NtPIP1;1t, NtTIP1;1s and NtNIP2;1s. The lowest H_2_O_2_ treatment concentration that resulted in a significant reduction in growth compared to untreated yeast was 0.25mM for NtPIP2;4s and NtPIP2;5t and 0.5mM for NtPIP1;1t, NtNIP2;1s and NtTIP1;1s.

**Figure 4.**
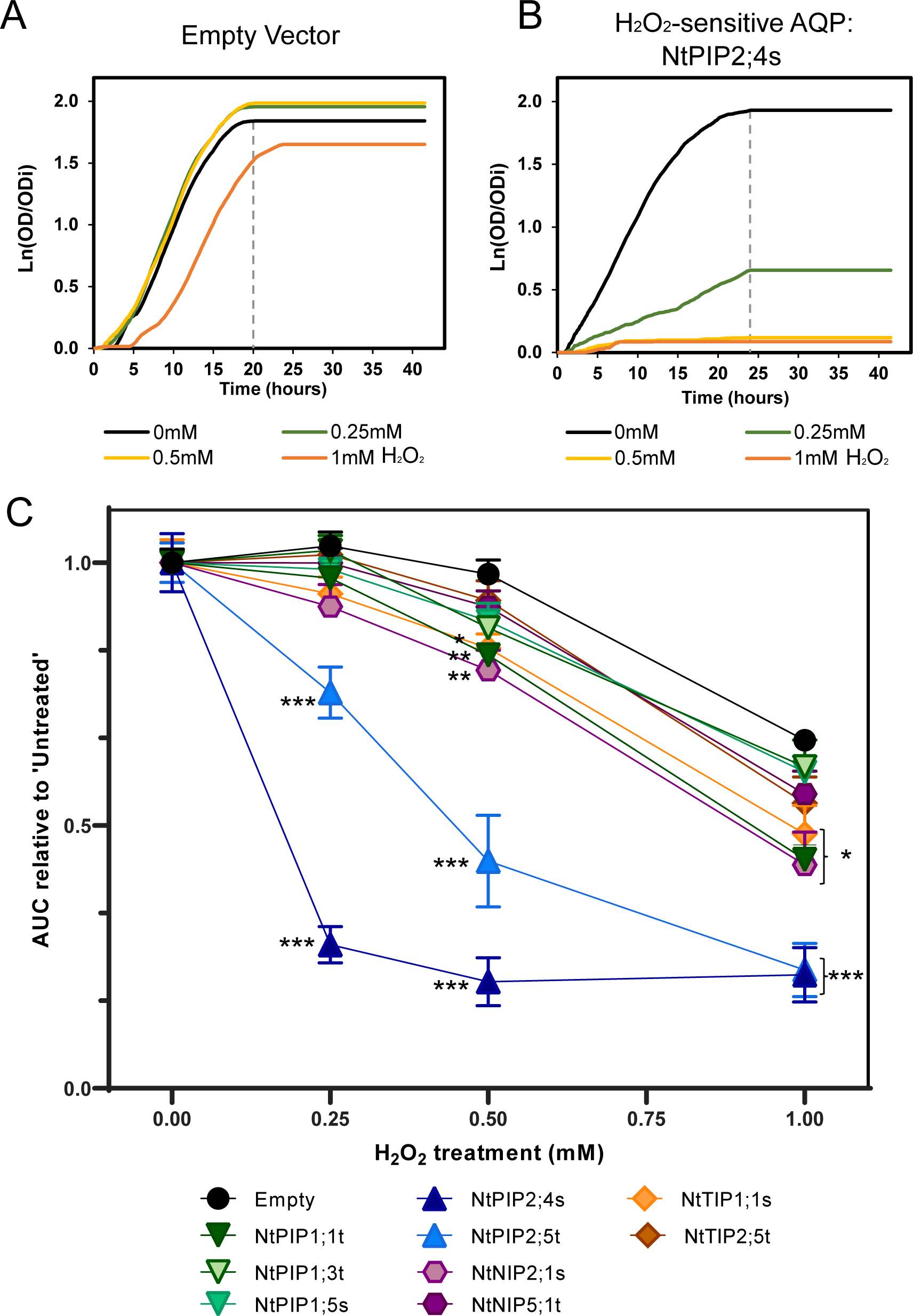
NtAQP H O toxicity screen. Yeast growth curves, Ln(OD/OD) vs. time, of snk7 yeast expressing **A.** Empty vector control or **B.** an H O -sensitive AQP (NtPIP2;4s), exposed to 0.25mM, 0.5mM and 1mM H O treatments. Growth was assessed from the area under the curves (AUC) until the vertical dashed lines. **C.** Yeast culture growth relative to ‘Untreated’ control (AUC relative to untreated), for skn7 yeast expressing an Empty vector or one the 9 NtAQPs. Asterisks denote One-Way ANOVA results comparing H O -treated yeast growth against Empty vector; *p<0.05, **p<0.01 and ***p<0.001. N=6, Error bars=SE.

### Boric acid toxicity assay

Toxicity-based assays were used to compare BA-sensitivity of yeast expressing the NtAQP isoforms and Empty vector control. Yeast expressing the empty vector (Figure 5A) showed reduced sensitivity compared to yeast expressing a “BA-sensitive” (NtTIP1;1s) AQP candidate, which had reduced growth in the presence of any of the BA treatments (10mM, 20mM and 20mM BA; Figure 5B). Exposure of yeast expressing the empty vector control to 10mM BA did not impair growth compared to untreated yeast (Figure 5A and 5C). However, greater BA concentrations progressively reduced growth (by 33% and 64% at 20 and 30mM BA, respectively). Three phenotypes were observed across NtAQP-expressing yeast exposed to 20mM and 30mM BA concentrations. The first phenotype displayed a severe BA-sensitivity phenotype, with a further 20-30% reduction in growth relative to the empty vector yeast (NtTIP1;1s and NtTIP2;5t, Figure 5C), consistent with these AQPs being classified as BA-permeable candidates, enhancing a toxicity response. The second phenotype had a BA toxicity response that was within 10-20% of the empty vector (NtNIP2;1s, NtNIP5;1t and NtPIP1;5s, Figure 5C). Within this phenotypic group, yeast expressing NtNIP2;1s had a significant reduction in growth at 10mM or 20mM BA (p<0.05), suggesting moderate sensitivity to BA exposure. The third phenotype, observed in 4 of the 5 PIP-expressing yeasts (NtPIP1;1t, NtPIP1;3t, NtPIP2;4s, NtPIP2;5t), resulted in a greater tolerance to BA exposure (average growth 20% greater than empty vector at 20mM and 30mM BA, Figure 5C). The reduced toxicity response associated with expressing any of these 4 NtPIPs could result from increased protein abundance in the PM reducing space available for free membrane diffusion of BA across the PM, thereby decreasing cell permeability to BA. Therefore, although NtPIP1;5s did not have a drastic decline in growth compared to the Empty vector control, growth was significantly reduced (∼25% at 20mM BA) compared to yeast expressing the other PIPs, likely due to facilitating boron transport across the PM.

**Figure 5.**
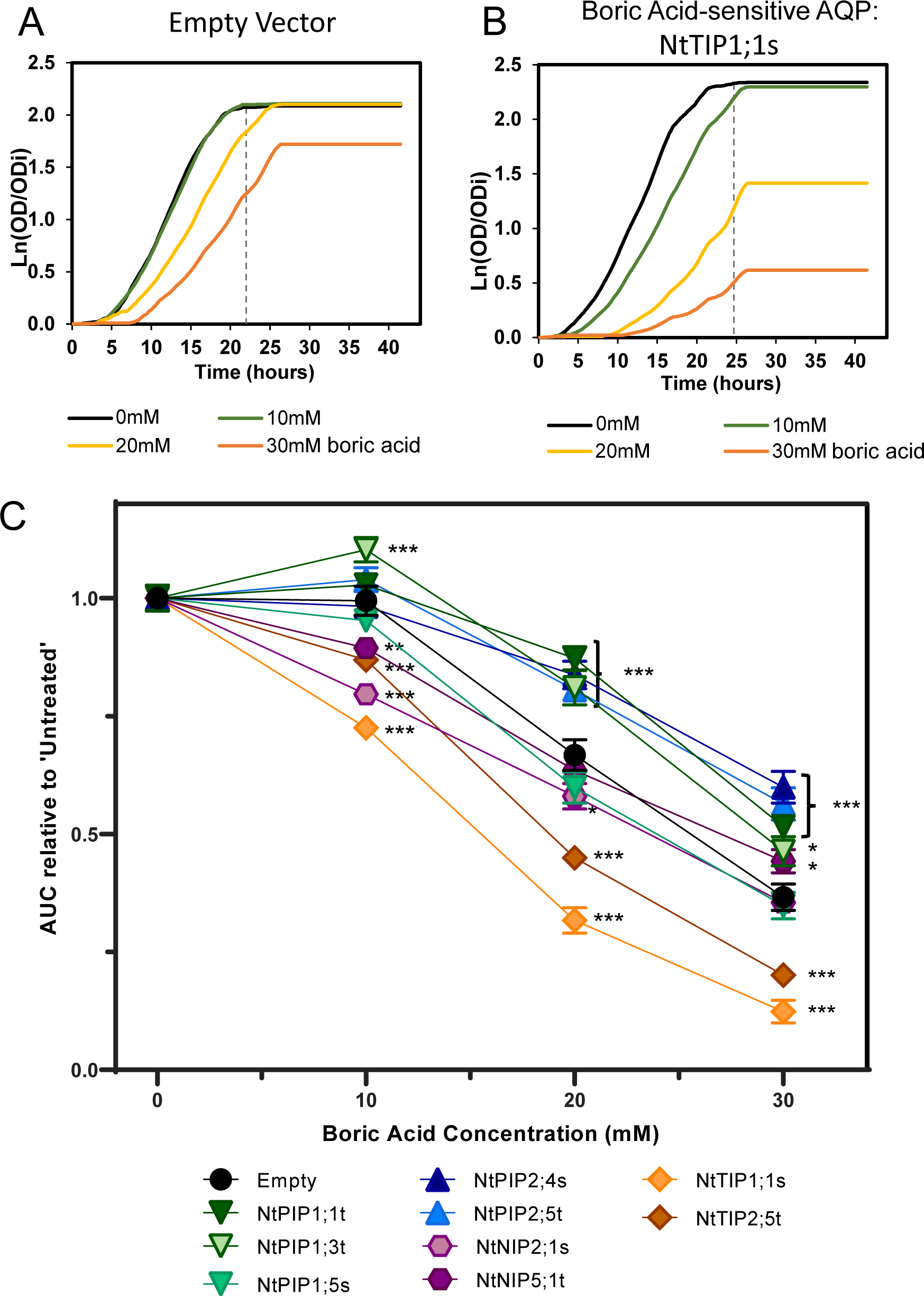
NtAQP boric acid toxicity screen. Yeast growth curves, Ln(OD/OD) vs. time, of *aqy1 aqy2* yeast expressing **A.** Empty vector control or **B.** a boric acid-sensitive AQP (NtTIP1;1s), exposed to 10mM, 20mM and 30mM boric acid treatments. Growth was assessed from the area under the curves (AUC) until the vertical dashed lines. **C.** Yeast culture growth relative to untreated control (AUC) of *aqy1 aqy2* yeast expressing either an Empty vector or one of the 9 NtAQPs exposed to boric acid. Asterisks denote growth that was significantly different to Empty Vector using a One-Way ANOVA; *p<0.05, **p<0.01 and ***p<0.001, N=6, Error bars=SE.

### Urea growth-based assay

Urea transport into yeast cells was tested through yeast complementation assays in the *ynvwI* strain lacking its native DUR3 urea transporter (Liu et al. 2003), where expression of urea-sensitive/permeable AQP candidates would result in enhancements of yeast growth phenotypes when exposed to low-urea/nitrogen source treatments. For *ynvwI* *y*east, 12mM urea provided sufficient nitrogen for yeast cultures to reach a growth curve plateau within a ∼50-hour incubation (Figure 6A-B, black lines). Growth curves of yeast expressing the empty vector (Figure 6A) or a “urea-sensitive” AQP (NtNIP2;1s, Figure 6B) show that the expression of the latter enhanced yeast growth at low urea concentrations (2mM and 4mM urea). Yeast expressing the empty vector exhibited a linear growth response to increasing urea concentrations (Figure 6C). Expression of NtTIP1;1s, NtNIP2;1s or NtTIP2;5t resulted in a 50% growth enhancement at both 2mM and 4mM urea compared to yeast expressing an empty vector (Figure 6C). Growth responses for the other 6 NtAQPs (NtPIP1;1t, NtPIP1;5s, NtPIP1;3t, NtPIP2;4s, NtPIP2;5t, NtNIP5;1t), were similar to the yeast expressing the empty vector at low urea concentrations (2mM and 4mM treatments) and these AQPs are presumed not permeable to urea.

**Figure 6.**
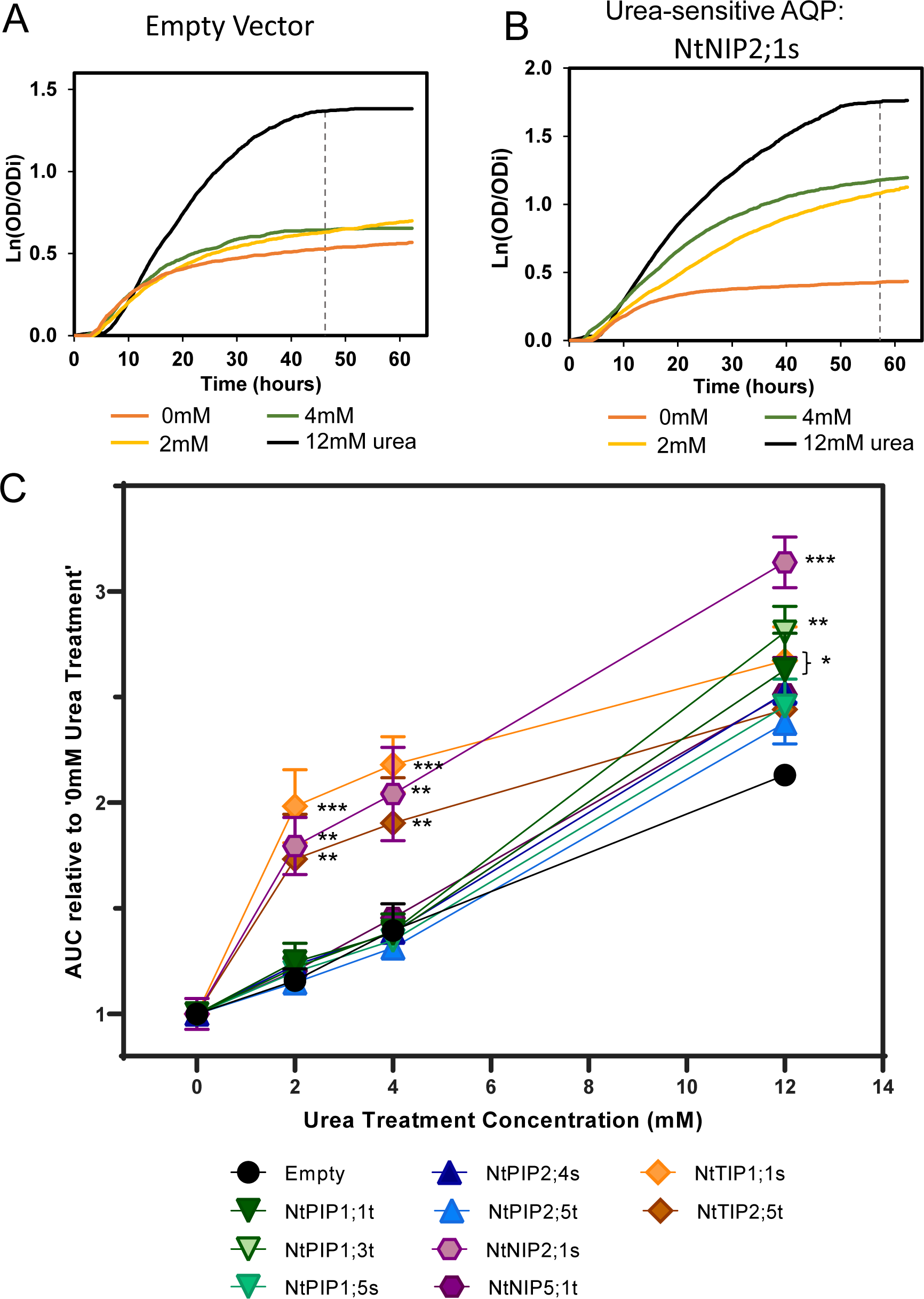
Urea growth-based assays for yeast expressing NtAQPs. Yeast growth curves, Ln(OD/OD) vs. time, of *ynvwI* yeast expressing **A.** Empty vector control or **B.** a urea-sensitive AQP (NtNIP2;1s), exposed to 0mM, 2mM, 4mM or 12mM urea treatments. Growth was assessed from the area under the curves (AUC) until the vertical dashed lines. **C.** AUC relative to 0mM Urea treatment of *ynvwI* yeast expressing either an Empty vector or one of the 9 screened NtAQPs. Asterisks denote growth that was significantly greater than Empty vector using a One-way ANOVA, *p<0.05, **p<0.01 and ***p<0.001. N=6, Error bars=SE.

### *In planta* sub-cellular localisation of tobacco AQPs

Arabidopsis transgenic lines over-expressing GFP-NtAQP or organelle specific marker lines were generated and confocal images of root cortical cells were captured (Figure 7). To enhance interpretation, surface plots of GFP intensity at magnified sub-regions near the cell wall were generated (Figure 7, white dashed box). Organelle-specific marker lines for the plasma membrane, ER and tonoplast and their distinct features were used as reference for interpretation of NtAQP subcellular localisation (Figure 7A-C). We observed substantial diversity in AQP membrane integration patterns across the NtPIP, NtTIP and NtNIP subfamilies. The NtPIPs showed a consistent signal around the periphery of the cell and a distinct absence of any internalised signal; a pattern similar to the PM marker. The NtPIP1s (NtPIP1;1t, NtPIP1;3t and NtPIP1;5s) appeared to have weaker and more diffuse PM integration compared to the sharp crips localisation of the NtPIP2s (PIP2;4s and PIP2;5t) (Figure 7D-H). The NtNIPs (NtNIP2;1s and NtNIP5;1t) also localised to the cell’s periphery. However, their GFP signal was speckled in appearance with distinct localised spots of brighter fluorescence characteristic of ER localisation (indicated by white arrow on NIP2;1s surface plot profile of GFP intensity, Figure 7I-J). The GFP signal of the NtNIPs was also broader than that of the PM-localising NtPIPs, consistent with colocalization to both the adjacent PM and ER. The localisation of NtTIPs (TIP1;1s and TIP2;5t) was consistent with integration in the tonoplast, showing a uniform yet diffuse localisation with a wavy topology; also denoted by the presence of internal membranes resembling transvacuolar strands (V, Figure 7K-L).

**Figure 7.**
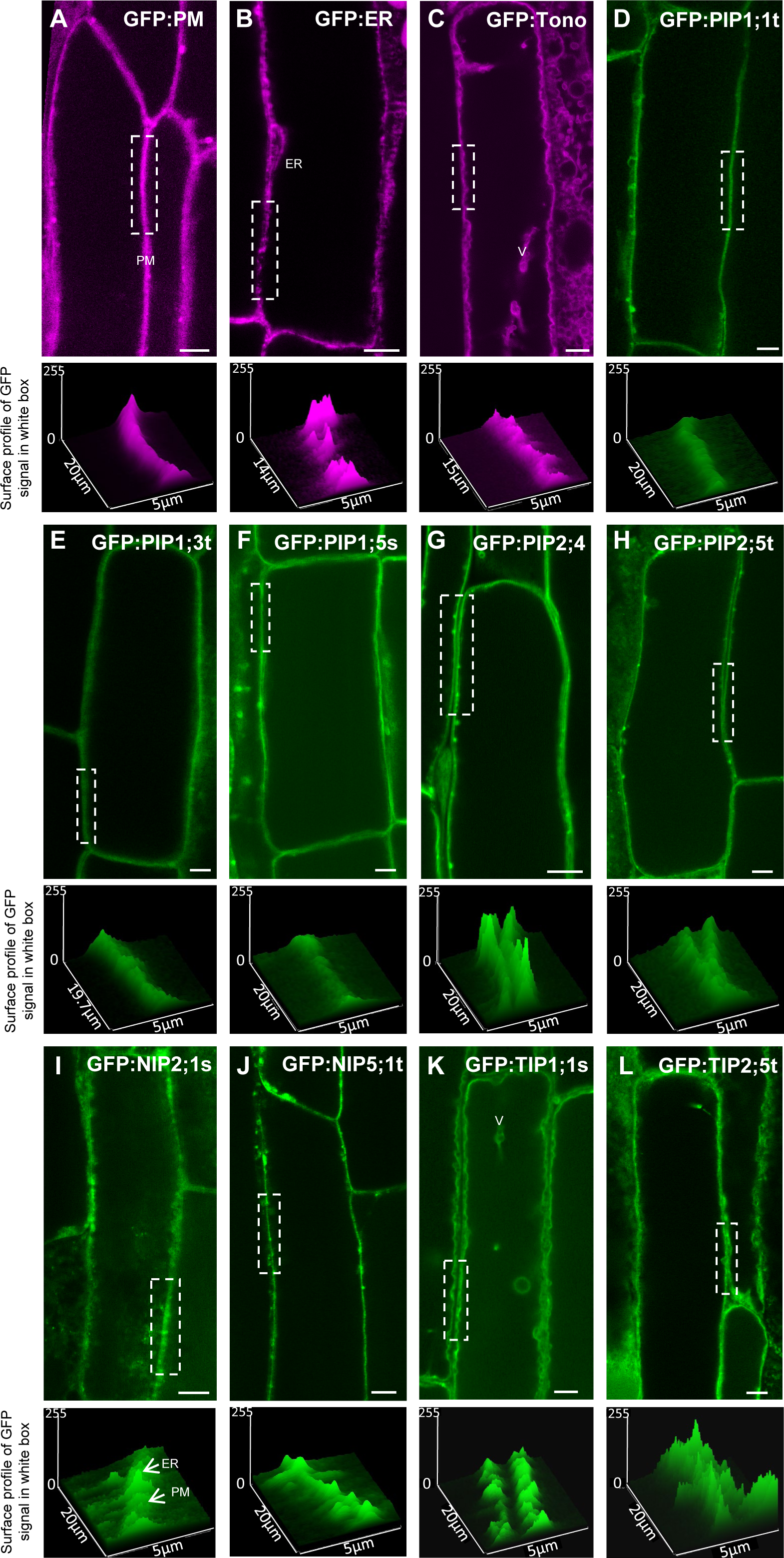
In planta sub-cellular localisation of NtAQPs. Confocal images of root cortical cells of transgenic 8-day old Arabidopsis seedlings. GFP marker lines; false coloured purple: **A.** plasma membrane, GFP:PM, **B.** endoplasmic reticulum, GFP:ER, **C.** tonoplast, GFP:tono. GFP:NtAQP lines: **D.** NtPIP1;1t, **E.** NtPIP1;3t, **F.** NtPIP1;5s, **G.** NtPIP2;4s, **H.** NtPIP2;5t, **I.** NtNIP2;1s, **J.** NtNIP5;1t, **K.** NtTIP1;1s and **L.** NtTIP2;5t. A region of the membrane (indicated by white dashed boxes, 5μm × 20μm dimension) is magnified in the panel below each confocal image to show surface profiles. Transvacuolar strands are denoted by **V.** White arrows highlight peak intensity discrepancies present in the NIPs assigned to AQP integration into the ER and PM. Scale bar 5μm.

### Protein modelling of NtAQP monomeric pores

Tertiary structure AlphaFold modelling and MoleOnline analyses were used to compare pore diameter and the physico-chemical properties (hydrophobicity and charge of pore-lining residues) of the 9 NtAQPs functionally characterised in this study (Figures 8-9). The generated pore diameter profiles of all PIPs closely overlapped (Supplementary Figure S1), as such NtPIP1;3t and NtPIP2;4s were used as representatives of NtPIP1 and NtPIP2, respectively, for less cluttered visual comparisons against NtTIP and NtNIP isoforms (Figure 8).

**Figure 8.**
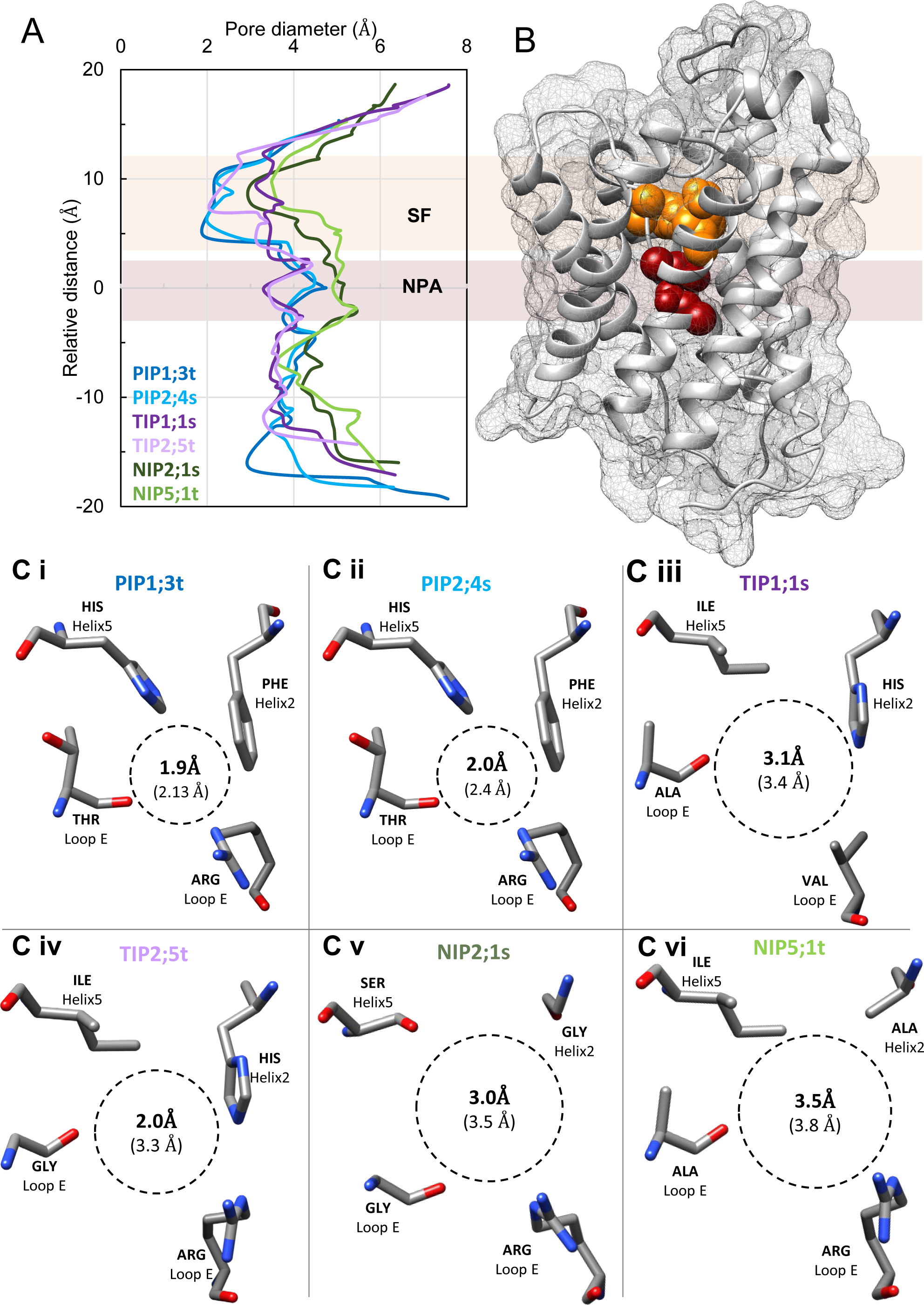
Modelled NtAQP pore features. **A.** Pore profiles of representative PIP1 (PIP1;3t, dark blue), PIP2 (PIP2;4s, light blue), TIP1;1s (dark purple), TIP2;5t (light purple), NIP2;1s (dark green) and NIP5;1t (light green). **B.** A 3D protein model highlighting the Selectivity Filter region (SF, orange residue in 3D Protein model) and NPA region (dark red residues in 3D protein model). **C.** Amino acid residues forming the selectivity filter (SF) and the diameter at its narrowest point in bold and the average SF diameter in brackets, viewed perpendicular to the membrane plane from the extracellular side for **C** i. PIP1;3, **C** ii. PIP2;4s, **C** iii. TIP1;1s, **C** iv. TIP2;5t, **C** v. NIP2;1s and **C** vi. NIP5;1t.

**Figure 9.**
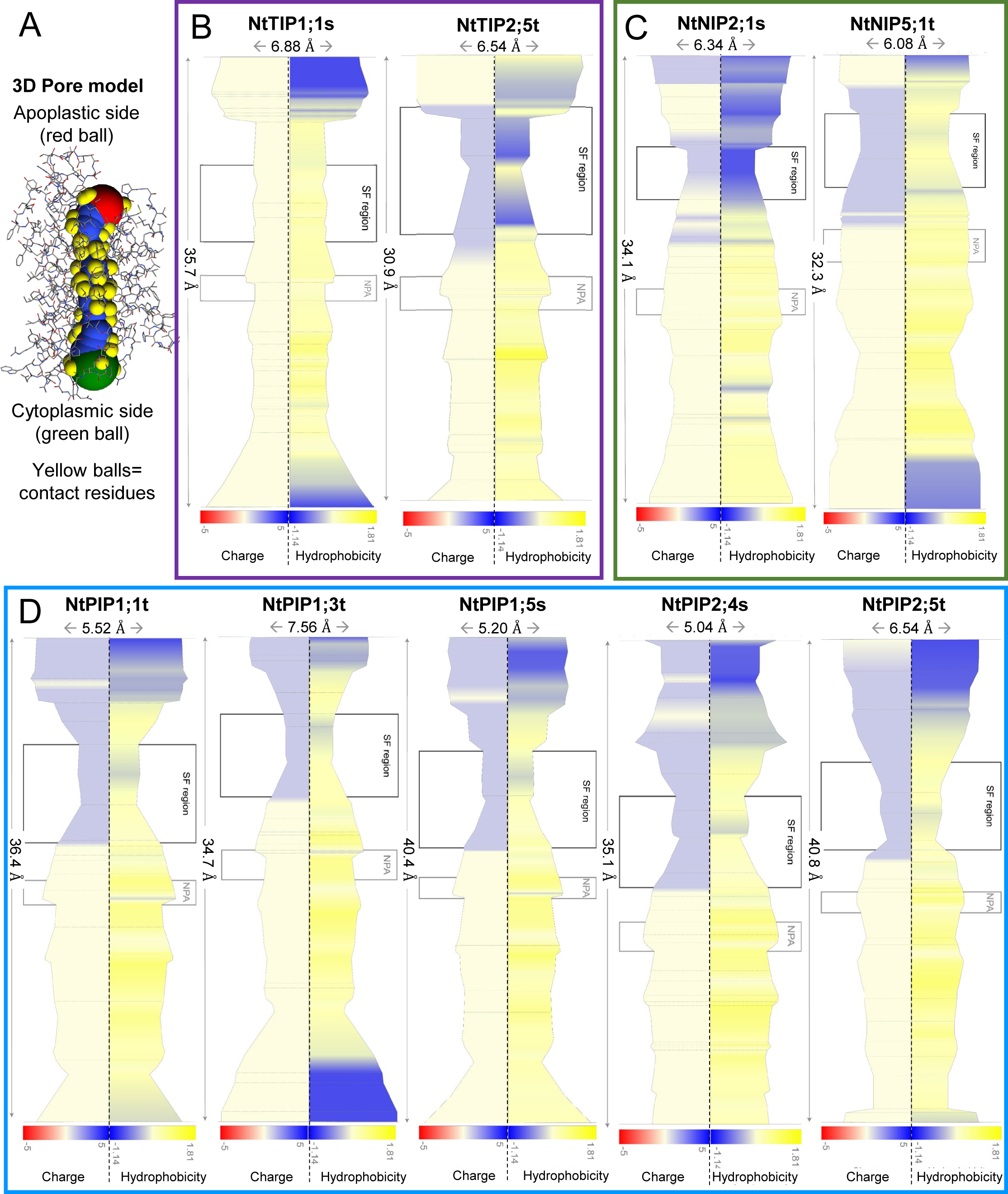
Hydrophobicity and charge profiles of NtAQP monomeric pores. **A.** 3D Pore model illustrates orientation of the pore profile with the apoplastic entrance (red ball, top), cytosolic entrance (green ball, bottom), residues contacting the pore (yellow balls) and the area inside the AQP pore (blue). Pore profile diameters from MoleOnline software (note maximum diameter and pore length scales for each AQP), of **B.** NtTIPs (NtTIP1;1s and NtTIP25t.) **C.** NtNIPs (NtNIP2;1s, NtNIP5;1t) and **D.** NtPIPs (NtPIP1;1t, NtPIP1;3t, NtPIP1;5s, NtPIP2;4s, NtPIP2;5t). Pore profiles are colour-filled with measures of “Charge” on the left (red to blue indicating positive i.e. ASP, LYS, HIS to negative i.e ASP, GLU, respectively) and “Hydrophobicity” (normalised hydrophobicity scale i.e. ILE =1.81 (Cid et al. 1992)) on the right (blue to yellow indicating low i.e. GLU = -1.14 to high hydrophobicity, respectively). Grey lines spanning the pore profiles correspond to contact residue interaction layers for which charge and hydrophobicity outputs were generated. Black boxes indicate Selectivity Filter (SF) region and grey boxes indicate NPA region.

Amongst the NtPIPs profiled, the Selectivity Filter (SF) region was the constriction point along the pore and the narrowest segment compared to all other NtAQP isoforms profiled (Figure 8A). The SF residue composition of the NtPIPs was highly conserved, each having Phe-His-Thr-Arg composition in Helix 2 (H2), Helix 5 (H5), Loop E position 1 (LE1) and Loop E position 2 (LE2), respectively. However, the PIP1s had a slightly narrower SF constriction (1.9Å vs 2.0Å) and average SF diameter (2.13Å vs 2.4Å) compared to the NtPIP2s (Figure 8c i-ii). The PIP monomeric pore physicochemical properties were also highly conserved, showing high similarity in residue charge and hydrophobicity (Figure 9D). All NtPIPs were lined with negatively-charged and hydrophilic residues (blue colours) from the apoplastic pore entrance to the SF region (Figure 9 D). All NtPIP1s also had a band of hydrophilic residues in the NPA region, however this was not present in the NtPIP2s. Additionally, NtPIP1;3t and to a lesser extent NtPIP1;1t and NtPIP2;5t were lined with hydrophilic residues in the cytoplasmic pore mouth. The remainder of the at PIP pore regions are lined with layers of hydrophobic residue bands starting from the SF region down to the cytoplasmic pore mouth (Figure 9D).

The NtTIPs (NtTIP1;1s and NtTIP2;5t) had broadly conserved monomeric pore diameter profiles (Figure 8A), and shared similarity with the monomeric pore profile of AtTIP2;1 published in (Kirscht et al. 2016) (Supplementary Figure S2). Notable differences were observed in the SF region between NtTIP2;5t and NtTIP1;1s, with NtTIP2;5t having a narrower SF constriction (2.0Å vs 3.1 Å), similar to the SF diameter of the PIPs, although the SF average diameter remained similar between the TIPs (3.3Å for NtTIP2;5t and 3.4 Å for NtTIP1;1s, Figure 8). The overall pore shape of the NtTIPs was nearly cylindrical/less undulating in comparison to the PIPs, with the SF region being wider for NtTIP1;1s and the NPA region being slightly narrower (3.3 Å in TIPs vs. the 3.8 Å of the PIPs; Figure 8A, Figure 9B). The pore properties profiles for NtTIPs differed between the two isoforms (Figure 9B). NtTIP1;1s had neutral charged residues throughput the pore and predominantly hydrophobic residues, with the exception of the pore entrances which was hydrophilic residues (Figure 9B). Alternatively, NtTIP2;5t had a pore profile similar to the PIPs, lined with negatively-charged residues in the SF region and hydrophilic residues in the apoplastic pore entrance and SF region (Figure 9B). The SF residue composition also differed between the 2 NtTIPs; with NtTIP1;1s having His- Ile-**Ala-Val** vs. NtTIP2;5t having His-Ile-**Gly-Arg** at H2-H5-LE1-LE2 positions (Figure 8C).

The NtNIPs pore diameter profiles were similar in shape and diameter (Figure 8A), and showed homology to the OsNIP2;1 monomeric pore (Saitoh et al. 2021) (Supplementary Figure S3). NtNIP2;1s and NtNIP5;1t have wide SF diameters, 3.5 Å and 3.8 Å average, respectively, a result of their SF composed of smaller residues than those found in other subfamilies (small residues bolded; NtNIP2;1s: **Gly-Ser-Gly**-Arg and NtNIP5;1t: **Ala**-Ile-**Ala**-Arg at H2-H5-LE1-LE2 positions respectively, Figure 8c). The NtNIPs had the widest NPA regions compared to NtPIPs and NtTIPs (4.9-5Å constriction, Figure 8). NtNIP2;1s had negatively charged and hydrophilic residues near the apoplastic pore entrance, the SF region and at various locations throughout the pore (Figure 9C). NtNIP5;1t had charged residues in the SF region and hydrophilic residues at both pore entrances and at banded regions in the SF (Figure 9C).

## Discussion

### Significance of determining NtAQP membrane localisation in yeast and *in planta*

In living systems, membranes are selectively permeable barriers enabling the separation of the cytoplasm from the cell exterior, and solute partitioning intracellularly within organelles (Gronnier et al. 2018; De Rosa et al. 2023). Selective transport of solutes across these membranes is crucial for cell functioning, with transport mechanisms (e.g. AQPs) providing capability to distinguish between chemically different solutes, and sense and respond to cellular transport and distribution requirements within the broad solute transport system (Stillwell 2016). As such, observing membrane localisation of AQP in cells is an important checkpoint for attributing solute transport at specific locations.

In yeast it is important to visualise PM localisation when undertaking heterologous expression assays, where the PM is the first barrier between the unicellular organism’s cytoplasm with external growth media solutions. We visualised AQP localisation in yeast using GFP translational fusions, confirming that all NtAQP isoforms integrated into the yeast PM (Figure 2), although to varying degrees. From these results we could then infer that variations in yeast growth phenotypes upon exposure to varying substrates could be due to AQP-related alterations in PM permeability (Bienert and Chaumont 2014). Although we observed PM-associated localisation for the NtPIP1s in yeast (Fig. 2), there are several cases where PIP1 permeability to water and/or other substrates has only been observed when PIP1 was co-expressed with a PIP2 in Xenopus oocytes, (Fetter et al. 2004) or weakly observed when the PIP1 was expressed alone and enhanced when co-expressed with a PIP2 in yeast expression systems (Groszmann et al. 2023), (reviewed in (Groszmann et al. 2017)). In our study, despite weak PM-integration of the NtPIP1 isoforms, we detected H_2_O_2_ and BA toxicity phenotypes for PIP1;1t and PIP1;5s, respectively. In future work, it would be useful to assess whether co-expression of NtPIP1s with NtPIP2s yields differing growth phenotypes to those observed in this study, with PIP1 expression alone.

In multicellular organisms such as plants, specific cell membrane localisations are required for distinct membrane transport functions: the PM holds pivotal importance in transport of solutes in and out of the cells as well as sensing cellular signals (Gronnier et al. 2018); the tonoplast membrane regulates the sequestration and retrieval of solutes from the vacuole for cell osmoregulation, as well as pH regulation, ion homeostasis, signal transduction and stress responses (Ozolina et al. 2022); and ER has the greatest surface area of all organelle membranes in plant cells and diverse functions, including protein and lipid production, signal transduction and cell-to-cell communication (Maeshima and Ishikawa 2008). AQP integration into a given membrane could imply a distinct role within the dynamic plant solute transport network (Figure 10). We observed diverse sub-family specific localisations in plant cell membranes, with PIPs localising to the PM, TIPs to the tonoplast and NIPs to the ER and PM (Figure 7, Figure 10). These broad sub-family localisation trends are consistent with those observed for AQP isoforms in other species (Wudick et al. 2009; Maeshima and Ishikawa 2008). AQP isoforms from the PIP and TIP subfamilies have also been localised to other cellular membranes such as the chloroplast envelope (Uehlein et al. 2008), although this was not assessed in this study. Further complexities in deciphering AQP localisation in planta include capturing the dynamics of AQP trafficking which alter their membrane localisation in their native species. Changes in AQP localisation can be triggered by AQP-AQP interactions, tissue specific or developmental responses and other biotic and abiotic stresses which might result in polarised integration into specific membranes (Yoshinari and Takano 2017; Ma and Yamaji 2015). Dynamic AQP localisation can also involve altered membrane targeting or internalisation from membranes, enabling swift changes to membrane solute permeability in response to cellular triggers (Chevalier and Chaumont 2015; McGaughey et al. 2018; De Rosa et al. 2023).

**Figure 10.**
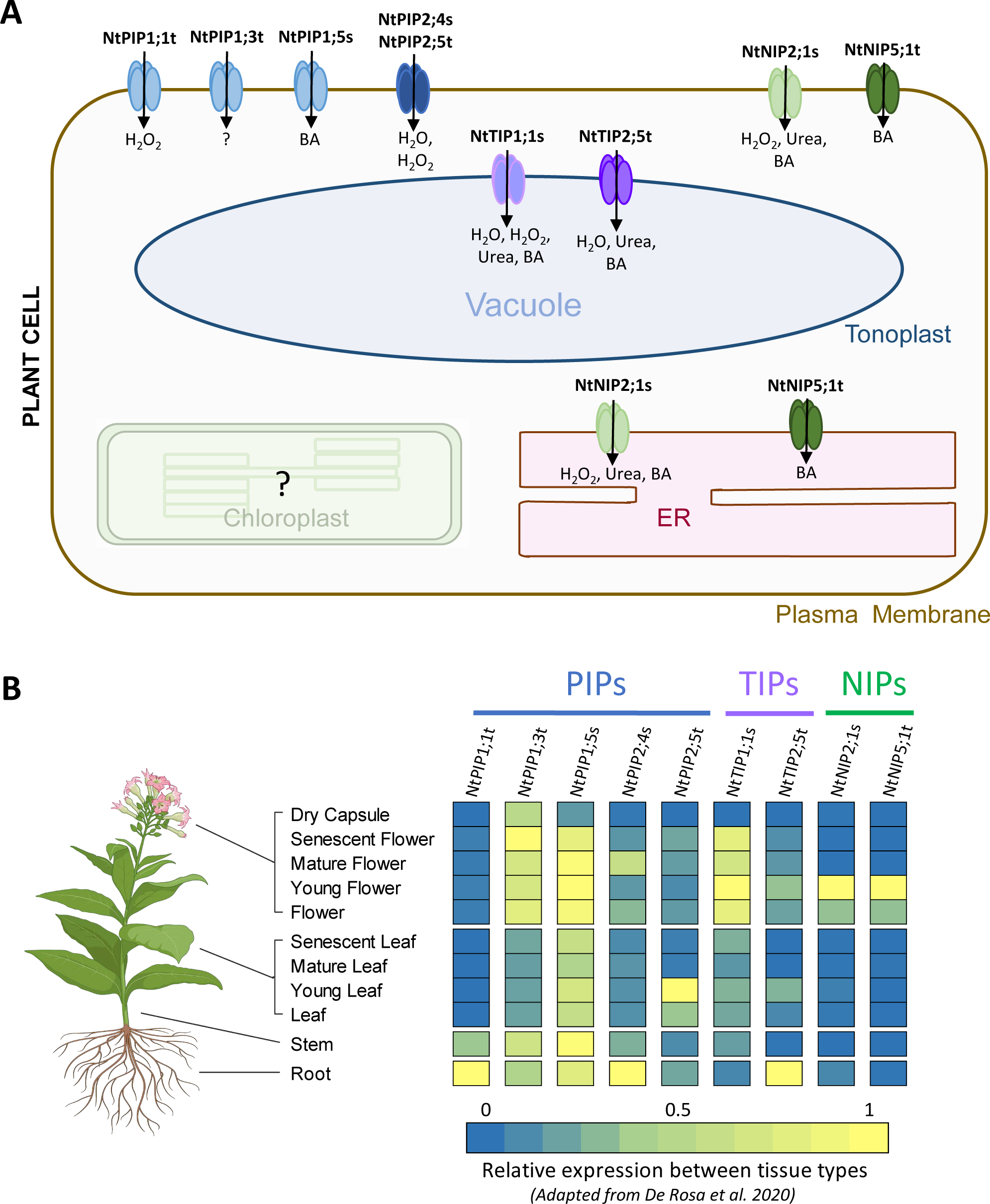
Summary of NtAQP putative solute transport, membrane localisation and gene expression profiles. **A.** Tobacco PIP1 (PIP1;1t, PIP1;3t and PIP1;5s, light blue), PIP2 (PIP2;4s and PIP2;5t, dark blue), TIP (TIP1;1s, light purple, and TIP2;5t, dark purple) and NIP (NIP2;1s light green, and NIP5;1t, dark green) isoforms screened in this study identified as novel candidates for water, H_2_O_2_, BA and urea transport, localised to the plasma membrane, tonoplast and ER in planta. **B.** Summary of Relative gene expression profiles of NtAQPs screened. Gene expression data adapted from (De Rosa et al. 2020). Tobacco Plant graphic created with BioRender.com.

### Relating functional characterisation, sub-cellular localisation, and gene expression of NtAQPs

We present the combined putative solute transport profiles, in planta sub-cellular localisation in Arabidopsis root cortical cells and gene expression profiles in tobacco for the 9 NtAQPs in Figure 10. In Table 1 the NtAQP data are further placed in context with other AQP candidates from other species with previously assigned solute permeabilities.

**Table 1.** Summary of NtAQP properties and other AQP isoforms characterised in other studies. Candidates tested for transport of water, H_2_O_2_, boric acid and urea; molecular diameter (Å) of substrates shown in parentheses as listed in (Azad et al. 2016); in planta sub-cellular localisation in root cortical cells with three sub-cellular localisations tested: plasma membrane (PM), endoplasmic reticulum (ER) and tonoplast; Selectivity filter (SF) constriction diameter and average SF region diameter in parentheses (Å), as identified from AlphaFold models; and gene expression localisations (reported in De Rosa et al. 2020). Tick (✓) denotes a positive assigned permeability for a specific substrate with the number of ticks indicating the magnitude of phenotypic responses observed. One, two and three ticks represent a small, medium and large phenotype effect, respectively, “C” denotes PIP1 identified as positive candidates when co-expressed with a PIP2. Grey shading of cells denotes a negative assigned/undetected permeability. Blank cells indicate no data is available.

|  | Water<br>(2.75 Å) | H<sub>2</sub>O<sub>2</sub><br>(3.2 Å) | Boron<br>(2.57 Å) | Urea<br>(2.62 Å) | In planta<br>subcellular<br>localisation | SF pore<br>diameter<br>(Å) | Expression<br>localisation | Ref. |
| --- | --- | --- | --- | --- | --- | --- | --- | --- |
| ZmPIP1 | C |  |  |  |  |  |  | (Fox et al. 2017) |
| AtPIP1;1 | C | ✓ | ✓✓ |  |  |  |  | (Groszmann et al. 2023) |
| NtPIP1;1t |  | ✓ |  |  | PM | 2.40 | Roots | This study |
| NtPIP1;3t |  |  |  |  | PM | 2.40 | Whole plant | This study |
| AtPIP1;4 | C | ✓✓ |  |  |  |  |  | (Groszmann et al. 2023) |
| NtPIP1;5s |  |  | ✓ |  | PM | 2.40 | Leaves, stem, roots, flowers | This study |
| AtPIP2;4 | ✓ | ✓ |  |  |  |  |  | (Groszmann et al. 2023) |
| NtPIP2;4s | ✓✓ | ✓✓✓ |  |  | PM | 2.40 | Roots, flowers (low) | This study |
| NtPIP2;5t | ✓✓ | ✓✓ |  |  | PM | 2.40 | Leaves | This study |
| ZmPIP2;5 | ✓✓ | ✓✓ |  |  |  |  | Roots, stem, leaves, | (Fox et al. 2017) |
| AtPIP2;7 | ✓✓ | ✓✓ | ✓ |  |  |  |  | (Groszmann et al. 2023) |
| ZmNIP1;1<br>NIP I | ✓✓ | ✓ | ✓ |  |  |  |  | (Fox et al. 2017) |
| AtNIP1;2<br>NIP I |  | ✓ |  |  |  |  |  | (Dynowski et al. 2008) |
| AtNIP2;1<br>NIP I | ✓ |  |  |  | ER |  | Young roots | (Mizutani et al. 2006) |
| AtNIP5;1<br>(NIP II) | ✓ |  | ✓✓ |  |  |  |  | (Takano et al. 2006; Wang et al. 2017) |
| NtNIP5;1t<br>NIP II |  |  | ✓ |  | ER + PM | 3.26 | Young flowers | This study |
| AtNIP6;1<br>NIP II |  |  |  | ✓✓ |  |  |  | (Wallace and Roberts 2005) |
| ZmNIP2;1<br>NIP III | ✓✓ | ✓ | ✓✓ | ✓✓ |  |  |  | (Fox et al. 2017) |
| NtNIP2;1s<br>NIP III |  | ✓ | ✓ | ✓✓ | ER + PM | 3.50 | Young flowers | This study |
| AtTIP1;1 | ✓✓ | ✓✓ |  |  |  |  |  | (Bienert et al. 2007) |
| NtTIP1;1s | ✓✓ | ✓ | ✓✓ | ✓ | Tonoplast | 2.75 | Leaves, flowers | This study |
| ZmTIP1;1 | ✓✓ | ✓✓ | ✓✓ | ✓✓ |  |  |  | (Fox et al. 2017) |
| AtTIP1;3 | ✓✓ |  |  | ✓✓ |  |  |  | (Soto et al. 2008) |
| AtTIP2;1 | ✓ |  |  | ✓ |  |  |  | (Liu et al. 2003) |
| **NtTIP2;5t** | ✓ |  | ✓✓ | ✓ | Tonoplast | 2.75 | Roots, leaves (low), flowers (low) | This study |

#### NtPIPs

The PIP subfamily generally has the largest number of isoforms (Anderberg et al. 2012), with PIPs implicated in broad solute transport roles relating to plant water homeostasis through highly selective water transport activity throughout the plant, as well as facilitating diffusion of other small molecules such as glycerol, urea, BA, arsenous acid, H_2_O_2_, gases and ions (Bienert et al. 2018; Liu et al. 2020). Certain isoforms are permeable to multiple solutes e.g. AtPIP2;1 permeable to water, H_2_O_2_, CO_2_ and cations, with functionality dictated by its location within the plant, interacting proteins and associated post-translational modifications (Tyerman et al. 2021). Such multifunctionality and regulation confers plant cells with broad solute transport capabilities throughout all tissues, responsive to cellular triggers. The 5 tobacco PIPs had high homology in SF composition (1.9-2 Å diameter), monomeric pore shape and physicochemical properties. However, our functional screens indicate diverse putative transport profiles between the NtPIP isoforms screened (Figure 10, Table 1). Therefore, other factors external to traditional specificity-determining features, such as other subtle sequence attributes, PIP1-PIP2 interactions, mechanisms impacting monomeric pore gating, and solute transport through tetrameric pore (e.g. for ions, larger solutes), could contribute to the broad solute transport capacity of PIPs (Tyerman et al. 2021).

Both NtPIP2s tested in this study were identified as water- and H_2_O_2_-permeable candidates, with yeast expressing NtPIP2;4s appearing more sensitive to H_2_O_2_ exposure than when expressing NtPIP2;5t. By contrast, none of the three NtPIP1s we screened were identified as water-transporting candidates. It is possible that co-expression of NtPIP1 with a NtPIP2 might enhance water transport across the yeast PM through improved PM localisation (Groszmann et al. 2023), but there was sufficient incorporation into the PM to enable detection of putative H_2_O_2_ (NtPIP1;1t) and BA (NtPIP1;5s) transport (Table 1). Similarly to the NtPIP solute transport profiles, several PIP isoforms from Arabidopsis (Groszmann et al. 2023), maize (Fox et al. 2017), rice (Kumar et al. 2014) and barley (Kumar et al. 2018) have been previously identified as H_2_O_2_ and/or BA permeable candidates, indicating functional conservation for the transport of these solutes from phylogenetically diverse PIPs. Further experiments could investigate whether PIP orthologs from closely related species, tomato and potato, which have similar gene expression profiles to the respective NtPIPs (De Rosa et al. 2020) also hold similar solute transport profiles. Specific signature sequence assessment by Ahmed et al. (2020) for tobacco AQPs, did not predict H_2_O_2_ transport for NtPIP1;1t nor BA transport for NtPIP1;5s, suggesting that subtle sequence differences or pore dynamics yet to be elucidated could impact preferential solute transport (Qiu et al. 2020). Other studies reported NtPIP1;5s having a low permeability to H_2_O_2_ using a fluorescence dye-based assay (Navarro-RóDenas et al. 2015). Since our yeast assay results did not show enhanced H_2_O_2_ sensitivity for NtPIP1;5s, this discrepancy needs to be confirmed using co-expression of PIP1 and PIP2 isoforms. Our screens did not identify any potential solute permeabilities for NtPIP1;3. This could be due to insufficient incorporation into the PM, or that NtPIP1;3t is permeable to substrates not tested here.

Integrating yeast assay functional data, subcellular localisation in planta and tissue specific expression analysis helps describe putative physiological roles for these NtPIPs within a broader solute transport system (Figure 10). The results implicate *NtPIP2;4s* and *NtPIP2;5t* having roles in regulating water transport across cell membranes in roots and leaves, respectively. Three of the NtPIPs tested here were permeable to H_2_O_2_, and likely to be involved with ROS signalling for a variety of developmental and stress cues, such as biotic and abiotic stress responses and plant developmental processes such as gemination (Hachez et al. 2006; Hoai et al. 2020) by facilitating H_2_O_2_ diffusion between cells. *NtPIP1;5s* (NtAQP1) was the first plant AQP shown to permeate CO_2_ (Uehlein et al. 2003), facilitating diffusion of CO_2_ into the chloroplast during photosynthesis (Flexas et al. 2006; Uehlein et al. 2008). We suggest *NtPIP1;5s* might have an additional role in boron uptake and distribution throughout the plant (expressed in roots, stems, leaves and flowers), similar to the functional roles reported for other boron-permeable PIPs, HvPIP1;3 and HvPIP1;4 (Fitzpatrick and Reid 2009) and OsPIP2;4 and OsPIP2;7 (Kumar et al. 2014).

#### NtTIPs

The TIP subfamily regulates the diffusion of water, ammonia, urea and metalloids across the tonoplast/vacuole membranes (Maurel et al. 1993; Loqué et al. 2005; Liu et al. 2003). Five specialised TIP subgroups have evolved in higher plants (TIP1-TIP5), differing in ar/R filter (SF) composition, substrate specificities (Anderberg et al. 2012; Kirscht et al. 2016), with some groups localising to the membranes of the central vacuoles (e.g. TIP1, TIP2 and TIP4), and others having been reported to localise to other vacuole-derived organelles (e.g. TIP3s localising to protein storage vacuole membranes) (Jiang et al. 2001; Maeshima and Ishikawa 2008). In addition to diversity in permeating substrates and vacuolar membrane localisation, the distinct TIP subgroups can further enable a staged control of vacuolar solute transport through facilitating transport at various developmental stages, e.g *TIP1s* are highly expressed during radicle elongation in germinating seeds and TIP2s are expressed after radicle emergence in fully elongated cells (Obroucheva 2013; Hoai et al. 2020).

Exemplars from both TIP1 (*NtTIP1;1s*) and TIP2 (*NtTIP2;5t*) subgroups localised to the tonoplast in planta, where they are implicated in transporting water, BA and urea (Figure 10). Yeast expressing NtTIP1;1s was also moderately sensitive to H_2_O_2_ but yeast expressing NtTIP2;5t were not. The substrate transport profiles we observed for NtTIP1;1s exactly match those for a Maize TIP1, ZmTIP1;1 (Fox et al. 2017), alluding to functional conservation of these TIP1-type isoforms.

The average SF region diameter of the NtTIPs was wider than that of the NtPIPs, although TIP2;5t had a similar SF constriction diameter to PIPs. An extended selectivity filter has been characterised for the TIP subfamily, containing an additional contact residue in Loop C of the AQP monomer (Kirscht et al. 2016), with NtTIP1;1s and NtTIP2;5t having a Phe and His at this position, respectively (De Rosa et al. 2020). The NtTIP1;1s SF composition of a Phe in Loop C and a Val in Loop E2 creates a more hydrophobic environment in the SF compared to that of NtTIP2;5t which has His and Arg in the same two positions respectively. This results in TIP1;1s being more hydrophobic than TIP2;5t in its SF region (Figure 9B).

The differences in SF composition and hydrophobicity between NtTIP1;1s and NtTIP2;5t might explain the divergence in permeability to H_2_O_2_ (Table 1) and potentially also for other predicted substrates such as ammonia (predicted for NtTIP2;5t but not NtTIP1;1s in Ahmed *et al*. (2020)).

Tissue-specific gene expression data revealed *NtTIP1;1s* is expressed in leaves and flowers whereas *NtTIP2;5t* is predominantly expressed in roots (having low expression in leaves and flowers). Root-specific gene expression of *TIP2* isoforms was also observed in closely related gene ortholog, tomato *TIP2;5* (De Rosa et al. 2020) and more distantly related maize *TIP2s* (Fox et al. 2017), indicating potential conservation of function of TIP2s across both closely related (tomato) and diverse (maize) species. The proposed functional roles for NtTIP1;1s and NtTIP2;5t in planta include the loading and unloading of urea from vacuolar storage, facilitating storage and translocation of boron, and equilibration of water homeostasis in the tissues where they are expressed (Maurel et al. 2015). Furthermore, NtTIP1;1s could also be involved in ROS signalling in response to cellular cues, as it has been suggested that TIPs are involved in cellular detoxification of H_2_O_2_ (Bienert and Chaumont 2014).

#### NtNIPs

NIP aquaporins are known to facilitate the transport of small uncharged solutes, such as glycerol, urea and metalloids (Wallace et al. 2006). NIPs generally have a more hydrophobic ar/R selectivity filter, which reduces water permeability in favour of other substrates such as ammonia, urea and metalloids (Wu and Beitz 2007; Hove and Bhave 2011). There are three NIP sub-classes (NIP I-III), based on ar/R selectivity filter and NPA motif composition (Mitani et al. 2008). Representatives from two of these sub-classes NIP II (*NtNIP5;1t*) and NIP III (*NtNIP2;1s*) were characterised here. GFP tagging demonstrated that both NtNIP2;1s and NtNIP5;1s were incorporated into the PM as well as localising to the ER in planta. NIP integration in the plant cell PM (versus tonoplast localisation of the TIPs) implies transport of these solutes in and out of cells, rather than storage/translocation from the vacuole which would be associated with TIPs (Figure 10).

NIP II aquaporins (such as NtNIP5;1t) tend to have a larger pore diameter than those found in the NIP I sub class, having a substitution of the highly conserved and bulky Trp at the ar/R H2 position for a smaller Ala (Wallace and Roberts 2004). NtNIP5;1t had a wider SF pore diameter (∼3.5 Å) compared to that of the tobacco PIP and TIP isoforms (Figure 8), and its pore diameter profile was also consistent with another recently modelled NIP II subclass isoform, AtNIP6;1 (Sharma et al. 2022). NIP II aquaporins have been shown to permeate BA, glycerol and urea, together with reduced water permeability compared to isoforms in the NIP I sub-class (Wallace et al. 2006; Takano et al. 2006; Hanaoka et al. 2014; Tanaka et al. 2008). While NtNIP5;1t was identified as a BA-permeable candidate, it was not identified to be sensitive to urea exposure, despite their similar size of these two substrates (Boron: 2.57 Å vs. Urea: 2.62 Å, Table 1). *NtNIP5;1t* expression is highly targeted to young flowers and could be involved in boron redistribution during flower development, similar to the orthologous gene in Arabidopsis (*AtNIP5;1*), which has an established role in boron transport and flower development (Takano et al. 2006). Notably, expression of NIP II isoforms, *AtNIP5;1* (in flowers) and *AtNIP6;1* (in basal shoot), is induced in boron-limiting conditions (Takano et al. 2006; Tanaka et al. 2008). The variation in expression patterns (localisation and stress-responsiveness) reported for PIPs vs. NIP IIs, allude to differences in physiological relevance of their boron transport in planta. PIPs could mediate a broad boron uptake and distribution (Fitzpatrick and Reid 2009), however their co-function as water channels subjects them to tight regulation (Chaumont and Tyerman 2014). Boron-permeable PIPs would also provide tolerance to boron toxicity by enabling efflux of excess boron from roots and shoots (Kumar et al. 2014; Kumar et al. 2018). Unlike the PIPs, NIP IIs tend to be poor water channels, enabling highly targeted boron transport in boron-limiting conditions (irrespective of water flux), and are down-regulated in boron sufficient concentrations (Figure 10) (Takano et al. 2006; Chaumont and Tyerman 2017).

NIP III aquaporins (such as NtNIP2;1s) are characterised by an ar/R filter composed of smaller residues (Gly-Ser-Gly-Arg), resulting in a wide, flexible and more hydrophilic SF region (Bansal and Sankararamakrishnan 2007; Mitani-Ueno et al. 2011). Pore diameter profile of a widely studied NIP III subclass representative, OsNIP2;1, shows an even wider SF constriction diameter than NtNIP2;1 (3.4-3.9Å, Supplementary Figure S3)(Saitoh et al. 2021). Although NtNIP2;1s’ SF constriction was not the widest of the NtNIPs screened (3.0 Å NIP2;1s vs. 3.5 Å NIP5;1t), its average SF diameter of 3.5 Å was nonetheless wider than other PIP and TIP isoforms screened (Figure 8), consistent with this sub-class being permeable to larger substrates, such as silicic acid and lactic acid (Mitani-Ueno et al. 2011). Unlike NtNIP5;1t (BA-permeable candidate only), NIP2;1s was identified as a candidate permeable to multiple substrates, urea, BA and H_2_O_2_ (low sensitivity), implicating this NIP isoform in multiple functional roles (Figure 10). NIP III isoforms occur widely among Graminae, but are not found in all dicots (e.g. absent in Arabidopsis), with evidence suggesting their principal role as facilitators of silicon uptake in plants (Chaumont and Tyerman 2017). In addition to silicon, ZmNIP2;1 (NIP III) is also permeable to water, urea, BA and H_2_O_2_ (low) (Fox et al. 2017), suggesting some functional homology in the maize and tobacco NIP III isoforms. Expression of *NtNIP2;1s* is restricted to young flowers where it is likely to be involved in strategic translocation of small molecules in this target tissue (Figure 10).

## Conclusion

Aquaporins are part of a dynamic solute transport network, spanning different tissues, developmental stages and cellular membranes. Improved understanding of AQP solute transport profiles, multifunctionality and structure-function relationships could help place the multitude of plant AQP isoforms within this network and elucidate putative key physiological roles.

This study characterised nine diverse isoforms in the tobacco AQP family for transport of key solutes: water, H_2_O_2_, urea and boric acid. The isoforms we chose localised to various cellular membranes in planta (PM, ER and tonoplast) and through yeast-based screens we identified candidates permeable to all, one or none of the substrates tested, indicative of broad or specific functional roles. The functional diversity observed between the NtAQP isoforms highlights complexity in assigning in planta function to specific isoforms, with monomeric pore shape/size, SF and NPA motifs alone insufficient to predict putative solute transport profiles.

## Methods

### Generation of NtAQP phylogeny

MUSCLE-aligned protein sequences from De Rosa et al. 2020 and Ahmed et al. 2020, (consensus NtAQP family in (Groszmann et al. 2021)), were used to construct a phylogenetic tree using neighbour-joining method (pair-wise deletion; bootstrap=1000) in MEGA7 software (Kumar et al. 2016).

### Generation of NtAQP expression constructs and transformation into yeast

Native sequences of *NtAQPs* sequences, *NtPIP1;1t* (BK011393), *NtPIP1;3t* (BK011396), *NtPIP1;5s* (BK011398), *NtPIP2;4s* (BK011406), *NtPIP2;5t* (BK011409), *NtTIP1;1s* (BK011426), *NtTIP2;5t* (BK011440), *NtNIP2;1s* (BK011379), *NtNIP5;1t* (BK011387) were commercially synthesised in Gateway-enabled destination vectors. Entry vectors were cloned into three destination vectors from (Alberti et al. 2007): pRS423-GPD with a Histidine3 (HIS3) marker gene for yeast selection, and pRS423-GPD-ccdB-ECFP and pRS426-GPD-eGFP-ccdB both containing Uracil3 (URA3) yeast selection gene. Yeast expression vectors were transformed in respective yeast strains required for functional assays (described below), using the “Frozen-EZ yeast Transformation Kit II” (Zymo Research, Los Angeles, USA). Transformed colonies were grown in Yeast Nitrogen Base, YNB, media (Standard drop out, DO, -URA or -HIS) and spotted on agar YNB (DO -URA or -HIS) selection plates for incubation at 30°C for 2 days, then stored at 4°C. Spotted plates were used for the starting cultures of functional assays.

### Confirming plasma membrane integration of NtAQPs in yeast cells

We assessed NtAQP subcellular localisations in yeast cells with AQP-GFP translational fusions to confirm incorporation of the expressed AQP in the yeast plasma membrane (PM). Tobacco AQP:GFP translational fusions were generated via Gateway cloning of pUC57 entry vectors with NtAQP coding sequences into pRS426-GPD-EGFP-ccdB yeast expression vector (Alberti et al. 2007); producing N-terminal GFP::NtAQP fusion proteins driven by the constitutive GPD promoter. The GFP-only yeast expression was obtained using the empty vector (no GOI fusion), with eGFP alone constitutively expressed via the GPD promoter. Yeast was grown overnight in 2mL YNB (-URA) (OD_600_ 1-1.5). 1mL aliquots of overnight cultures were sub-cultured and grown 3-4 hours in 2mL of fresh YNB (-URA) media, to ensure imaging of newly formed cells. Yeast (10μL) was mounted on a polysine slide with a coverslip sealed with nail polish. Yeast cells were visualised with a Zeiss LSM 780 Confocal microscope using a 40x oil immersion objective (1.2 NA). Light micrographs of yeast cells were acquired using Differential Interference Contrast (DIC), with GFP fluorescence captured using excitation at 488 nm and emission detection across the 490-526 nm range. Images were processed using Fiji (ImageJ) software (Schindelin et al. 2012).

### Assessing water, H_2_O_2_, boric acid and urea permeability using high-throughput yeast-based assays

High-throughput yeast microculture assays described in Groszmann et. al (2023) were used to screen NtAQPs for transport of specific substrates, associating yeast growth enhancements or toxicity phenotypes to expression of foreign AQPs in yeast cells in response to various treatments. Yeast growth was monitored using a SPECTROStar nano absorbance microplate reader (BMG Labtech, Germany) at 10-20 minute intervals over 42 to 60 hours. Data collection and processing was consistent between each growth or toxicity-based assay.

#### Water permeability “Freeze-thaw” assay

We used the *aqy1 aqy2* yeast strain (null aqy1 aqy2; background Σ1278b; genotype: Mat α; leu2::hisG; trp1::hisG, his3::hisG; ura352 aqy1D::KanMX aqy2D::KanMX, provided by Peter Dahl of the S. Hohmann lab) (Tanghe et al. 2002) exploiting the property of yeast cells that show increased freezing tolerance when they express functional water-transporting AQPs (Deshmukh et al. 2016). Yeast expressing NtAQPs and the empty vector were grown for 24-28 hours (OD_650_ of 0.5-1) in 1.25mL YNB(-HIS), at 30°C, shaking at 250rpm. Cultures were diluted to 0.6×10^7^ cells/mL in YPD medium and incubated at 30°C for 60 mins. 250μL of each culture was aliquoted to 2 Eppendorf tubes; one tube was placed on ice (untreated control) the other was used for two ‘Freeze-thaw’ cycles. Each ‘Freeze thaw’ cycle consisted of yeast culture aliquots being frozen in liquid nitrogen for 30 seconds, and thawed in a water bath at 30°C for 20 mins. For each construct, ‘Untreated’ and ‘Treated’ yeast were transferred into a 96 well plate (200μL aliquots) for growth monitoring.

#### H_2_O_2_ toxicity assay

Toxicity phenotypes of AQP-expressing yeast was assessed using a reactive oxygen species (ROS) hypersensitive yeast strain, *Δskn7* (null skn7; background BY4741 genotype: Mat α; his3Δ1 leu2Δ0 met15Δ0 ura3Δ0 ΔSKN7), obtained from ATCC (Bienert et al. 2007; Halliwell and Gutteridge 2015; Lee et al. 1999). Yeast’s survival was further compromised if AQPs facilitated the diffusion (and accumulation) of H_2_O_2_ into the cell, which enhanced the toxicity response. *Δ*skn7 yeast expressing NtAQPs and Empty vector were grown and diluted to 0.6×10^7^ cells/mL as per the Freeze-thaw assay (above). 200μL microcultures of each NtAQP/Empty vector were distributed in 96-well plates with 190μL of yeast and 10μL H_2_O_2_ treatments: 0mM/water, 0.25mM, 0.5mM and 1mM H_2_O_2_.

#### Boric acid (BA) toxicity assay

was used to screen for enhanced AQP-associated sensitivity of yeast cells upon exposure to increasing BA treatment concentrations. *aqy1 aqy2* yeast expressing NtAQPs or Empty vector were grown and diluted to 0.6×10^7^ cells/mL. 200μL microcultures each NtAQP/Empty vector were distributed in 96-well plates with 180μL of yeast and 20μL BA treatments: 0mM/water, 10 mM, 20 mM and 30 mM BA.

#### Urea growth-based assay

*ynvwI* yeast (null dur3; background Σ23346c; genotype: Mat α, Δura3, Δdur3, provided by Patrick Bienert of the Nicolaus von Wirén lab) is limited in growth due to a deletion of the DUR3 urea transporter (Liu et al. 2003). Expression of urea-permeable AQPs candidates in *ynvwI* yeast provided a growth advantage when exposed to media containing urea as the sole nitrogen source. *ynvwI* yeast spots were resuspended in 1.25mL of Yeast Basic media (YB, culture medium without nitrogen source) with 2% Glucose. Yeast cultures were diluted to 1.2×10^7^ cells/mL. 200μL microcultures for each NtAQP/Empty vector construct were distributed in 96-well plates with 190μL of yeast and 10μL urea treatments: 0mM/water, 1 mM, 4 mM and 12 mM urea.

#### Data Processing

The collected yeast microculture OD_650_ readings for the Freeze-thaw (water), H_2_O_2_, BA and urea transport assays were processed as described in Groszmann et al. (2023). Briefly, yeast growth was measured over time with yeast growth curves represented as natural log of OD_650_/initial OD_650_ (Ln(OD_t_/OD_i_) vs time; and a biologically meaningful measuring time point (Phi ф, when growth had fallen to 5% of the maximum growth rate of ‘Untreated’ yeast culture) was calculated for each yeast culture expressing NtAQPs and the Empty vector and across treatments. This enabled standardisation of growth measurements and comparison of growth of yeast expressing various constructs, each exposed to different treatments. Area Under the Curve (AUC) was obtained from the yeast growth curves, as a proxy for yeast growth providing a cumulative measure of growth, capturing growth characteristics occurring during both the lag and growth phases of the yeast culture, enabling us to capture subtle cumulative growth effects resulting from the treatments (Groszmann et al. 2023).

### Characterising *in planta* subcellular localisation of NtAQPs

Tobacco AQP-GFP constructs were generated via Gateway cloning of NtAQP coding sequences from pZeo entry vectors into the pMDC43 destination vector (Curtis and Grossniklaus 2003); N-terminal GFP-NtAQP fusion proteins were driven by the constitutive 2×35S CaMV promoter. Arabidopsis transgenic lines were generated via agrobacterium (GV3101) floral dipping transformation method (Clough and Bent 1998). Seeds were liquid sterilised, washed and sown on Gamorg’s B5 medium (0.8% Agar, hygromycin). 8-day-old Arabidopsis seedlings were gently removed from the agar, mounted in phosphate Buffer (100mM NaPO_4_ buffer, *pH* 7.2) on a standard slide, covered with coverslip, and visualised with a Zeiss LSM 780 Confocal microscope using a 40x water immersion objective (1.2 NA). Light micrographs of cortical cells in the root elongation zone were visualised using Differential Interference Contrast (DIC), GFP fluorescence was captured using excitation at 488 nm and emission detection across 490-526 nm. Autofluorescence was detected across 570-674 nm and excluded from GFP detection channel. Images were processed using Fiji (ImageJ) software (Schindelin et al. 2012). Organelle-specific marker lines, established in Nelson et al. (2007) and previously published in De Rosa et al. (2020), were used to guide our interpretation of AQP subcellular localisations.

### 3D AlphaFold protein modelling and characterisation of NtAQP pores

The peptide sequences for the 9 NtAQPs characterised in this study were used to generate 3D protein models through artificial intelligence prediction methods. These methods incorporate known physical and biological properties of protein structure to predict protein folding conformations (Jumper et al. 2021). We modelled all 9 NtAQP structures using ColabFold, an open-source webserver based on AlphaFold2 (Mirdita et al. 2022). Output models (PDBs) obtained from ColabFold were ranked by predicted local distance difference test (plDDT) scores, where higher scores indicate greater stereochemical plausibility of a model (Mariani et al. 2013). The model with highest pIDDT score was selected for monomeric pore characterisation analyses. PDB files of NtAQP models were uploaded to MOLE*online* webserver (https://mole.upol.cz/; Web UI Version: 1.5.2.1), an online interface to locate and characterise channels, pores, and tunnels in macromolecular structures (Pravda et al. 2018). We computed the most probable monomeric pore of all NtAQPs; input settings, pore coordinates and pore-lining contact residues obtained for each NtAQP isoforms are listed in Supplementary File 1. The selectivity filters and asparagine-proline-alanine (NPA) regions characterised in De Rosa et al. 2020 were mapped along the pore profile/pore lining contact residues for each NtAQP. For ease of comparison of monomeric pore profiles across isoforms, the ‘relative’ distance was standardised to the first contact point (from the cytosolic side) of the respective ‘asparagine’ for the two NPA motifs.

## Acknowledgements

We thank Rosemary White from CSIRO for providing seeds of the Tonoplast:GFP and ER:GFP marker lines (Nelson et al. 2007). The authors acknowledge the facilities and the scientific and technical assistance (Darryl Webb) of Microscopy Australia at the Advanced Imaging Precinct at the Australian National University; a facility funded by the ANU, and State and Federal Governments of Australia. We acknowledge that National Collaborative Research Infrastructure Strategy (NCRIS) of the Australian Government, providing The Australian National University with the growth facilities utilised as part of the Australian Plant Phenomics Facility.

## Sentence Summary

Diverse tobacco PIP, TIP and NIP aquaporin isoforms were functionally characterised using high-throughput yeast-based assays, identifying candidate transporters for key plant solutes: water, H_2_O_2_, boric acid and urea.

## Author Contributions

ADR, MG and JRE conceived research plans; ADR performed yeast screening experiments, subcellular localisation analyses and wrote the manuscript; ADR and RZ performed 3D protein modeling analyses; ADR, RZ, JRE and MG analyzed the data. ADR, CB, JRE and MG critically reviewed and edited the manuscript. ADR agrees to serve as the author responsible for contact and ensures communication.

