## Supplementary material for "Exploring the Plant Aquaporin Solute Transport Network: Functional characterisation of *Nicotiana tabacum* PIP, TIP and NIP isoforms": Suplementary Figures

### Supplementary Data

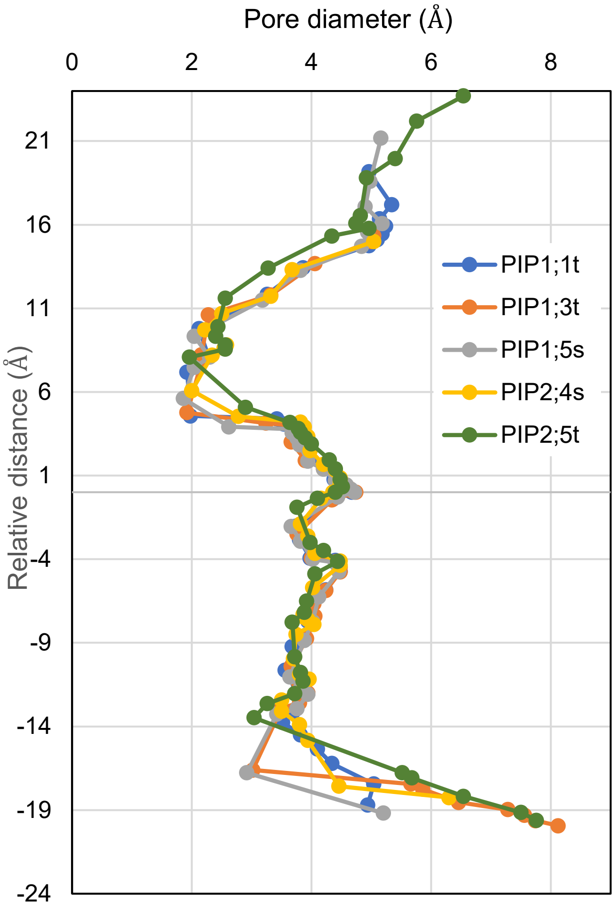


**<u>Supp Figure 1.</u>**
**Pore Diameter profiles of all NtPIPs.**

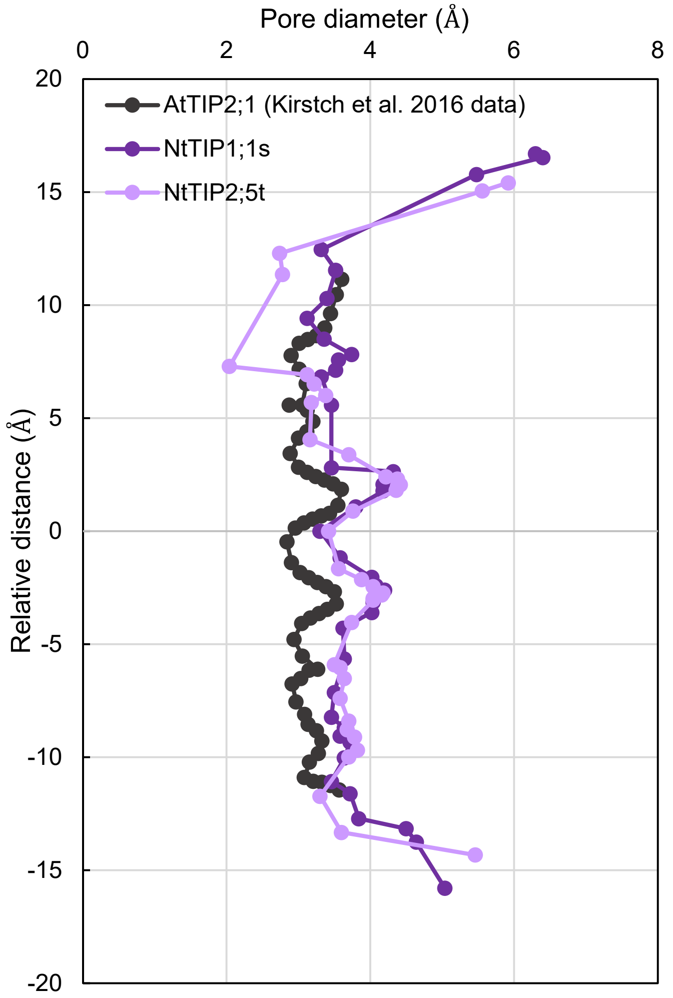


**<u>Supp Figure 2.</u> Comparison of Pore diameter profiles of NtTIPs (NtTIP1;1s and NtTIP2;5t) and Arabidopsis TIP2;1 (Kirstch et al. 2016).**

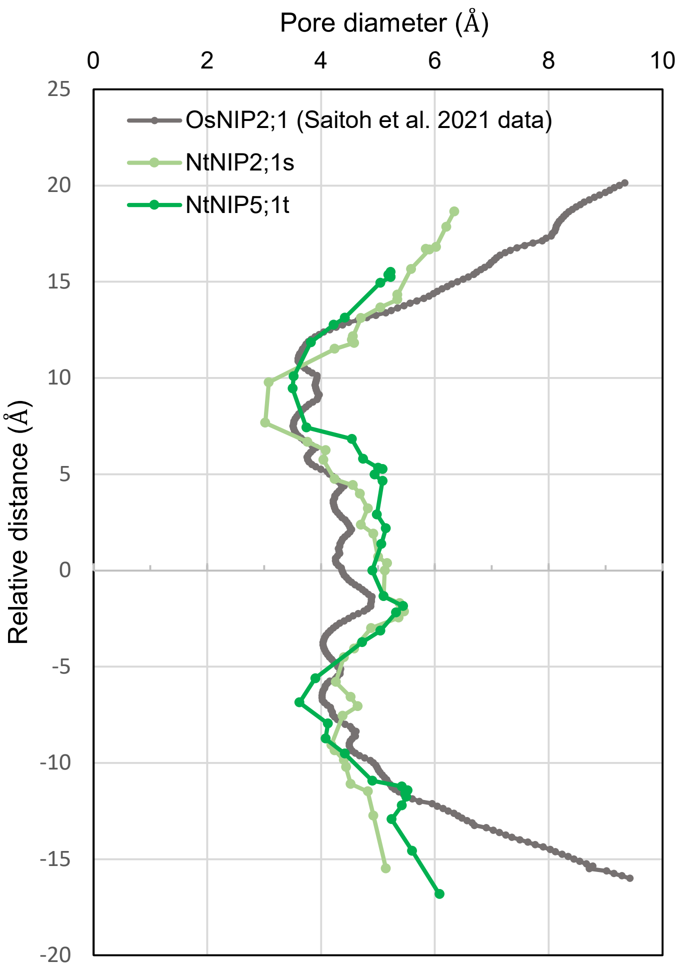


**<u>Supp Figure 3.</u> Comparison of Pore diameter profiles of NtNIPs (NtNIP2;1s and NtNIP2;5t) and Oryza sativa NIP2;1 (Saitoh et al. 2021).**
